# Dual Role of Jam3b in Early Hematopoietic and Vascular Development

**DOI:** 10.1101/656108

**Authors:** Isao Kobayashi, Jingjing Kobayashi-Sun, Yuto Hirakawa, Madoka Ouchi, Koyuki Yasuda, Hiroyasu Kamei, Shigetomo Fukuhara, Masaaki Yamaguchi

## Abstract

In order to efficiently derive hematopoietic stem cells (HSCs) from pluripotent precursors, it is crucial to understand how mesodermal cells acquire hematopoietic or endothelial identity due to their close developmental connection. Although Npas4 has been recently identified as a conserved master regulator of hemato-vascular development, the molecular mechanisms underlying the cell fate divergence between hematopoietic and vascular endothelial cells are still unclear. Here, we show in zebrafish that the divergence of hematopoietic and vascular endothelial cells in mesodermal cells is regulated by Junctional adhesion molecule 3b (Jam3b) via two independent signaling pathways. Mutation of *jam3b* led to the reduction of *npas4l* expression in the posterior lateral plate mesoderm and defect of both hematopoietic and vascular development. Mechanistically, we uncover that Jam3b promotes endothelial specification by regulating *npas4l* expression through the repression of the Rap1a-Erk signaling cascade. Jam3b subsequently promotes hematopoietic development including HSCs by regulating *lrrc15* expression in endothelial precursors through the activation of an integrin-dependent signaling cascade. Our data provide insight into the divergent mechanisms for instructing hematopoietic or vascular fates from mesodermal cells.

## Introduction

Hematopoietic stem cells (HSCs) can produce all types of mature blood cells throughout the lifespan of an individual and are the therapeutic component of bone marrow transplantation. One of the major goals of regenerative medicine is to derive HSCs from induced pluripotent stem cells (iPSCs). It is of great interest to better understand the developmental mechanisms of HSCs in the embryo because elucidation of this process can provide insight into the cellular and molecular cues needed to instruct and expand HSCs from iPSCs (Murry and Keller, 2008). In vertebrates, HSCs arise from a subset of endothelial cells, termed hemogenic endothelial cells, in the ventral floor of the dorsal aorta (DA) (Boisset et al., 2010; Bertrand et al., 2010; Kissa and Herbomel, 2010). Studies in mouse and zebrafish embryos have greatly contributed to our understanding of the cellular and molecular programs controlling HSC development. Some transcription factors, such as Runx1, c-Myb, Scl (also known as Tal1), and Gata2, have been implicated directly in the regulation of HSC formation and maintenance during embryogenesis (Clements and Traver, 2013). It is believed that hematopoietic identity is conferred through the expression of these transcription factors within endothelial precursors. However, the molecular mechanisms underlying the hematopoietic specification is still largely unknown, leading to difficulties in efficient derivation of hematopoietic lineages from iPSCs.

Hematopoietic and vascular endothelial cells develop in close proximity to each other within the mesoderm. It has been proposed that these two cell populations share a common precursor, termed the “hemangioblast” (Choi et al., 1998; Huber et al., 2004; Vogeli et al., 2006). The zebrafish *cloche* mutant, in which both hematopoietic and vascular development are defective, has been identified as a unique model for the study of hemangioblast development (Stainier et al., 1995). Due to the reduction of various endothelial and hematopoietic marker genes in this mutant embryo, the *cloche* gene is thought to act at the top of hierarchy for hemangioblast specification (Sumanas et al., 2005). Recently, it has been determined that the *cloche* gene encodes a PAS-domain-containing bHLH transcription factor, Npas4l (neuronal PAS domain protein 4 like). Importantly, enforced expression of its human orthologue, *NPAS4*, can rescue the vascular defect in *cloche* mutant embryos, suggesting that NPAS4 plays a central role in the hemato-vascular development in an evolutionarily conserved manner (Reischauer et al., 2016).

Junctional adhesion molecule (Jam) proteins belong to the immunoglobulin (Ig) superfamily cell adhesion molecule and consist of two Ig-like domains, a transmembrane domain, and a type II PDZ-binding motif. Jam3 is widely expressed on the epithelium, endothelium, and some leukocytes, and has been implicated in tight junction formation and barrier function as well as leukocyte migration (Ebnet et al., 2017). Heterophilic interactions between Jam2 (also known as Jam-B) and Jam3 play an important role in leukocyte transmigration (Weber et al., 2007), spermatozoal differentiation (Gliki et al., 2004), and myocyte fusion (Powell et al., 2011). In mice, Jam3 is expressed on adult HSCs and promotes the adhesion of HSCs to Jam2-expressing stromal cells in the bone marrow (Arcangeli et al., 2011). Inhibition of Jam3 by a blocking antibody affected the homing and reconstitution ability of hematopoietic stem/progenitor cells (HSPCs) (Arcangeli et al., 2014). Jam3 also mediates an increase in endothelial permeability through repression of the Rap1 activity, a member of the Ras family of GTPases that can enhance the adhesive contact between adjacent endothelial cells (Orlova et al., 2006). Although Jam3 functions in both hematopoietic and vascular endothelial cells, to date, there has been no reported role for Jam3 in the development of hematopoietic or vascular endothelial cells.

Here, we show that a zebrafish orthologue of Jam3, Jam3b, plays dual roles in hemato-vascular development during early embryogenesis: (1) it promotes endothelial specification in mesodermal cells via the regulation of *npas4l*, and (2) it confers hematopoietic fate in endothelial precursors through the regulation of Leucine rich-repeat containing 15 (Lrrc15). In vertebrates, the mesodermal induction requires fibroblast growth factor (FGF)-dependent extracellular signal-regulated kinase (Erk) signaling (Umbhauer et al., 1995; Kimelman, 2006; Xiong et al., 2015), a component of the mitogen-activated protein kinase (MAPK) cascade. We found that Jam3b suppresses the activation of Erk to enhance *npas4l* expression in the lateral plate mesoderm (LPM). Jam3b subsequently induces *lrrc15* expression in endothelial precursors through an integrin-dependent signaling cascade, driving them to acquire the HSC fate.

## Results

### Jam3b is required for both hematopoietic and vascular development

To investigate if *jam3b* is expressed in hematopoietic and vascular endothelial cells in zebrafish embryos, we performed whole-mount *in situ* hybridization (WISH) using a *jam3b*-specific probe. The expression of *jam3b* was first detected at around 8 hours post-fertilization (hpf) mainly in the dorsal part of the embryo, while it was undetectable by 6 hpf (Fig. S1A). At 14 hpf, we could detect *jam3b* expression in the posterior LPM (PLPM) where hematopoietic and endothelial precursors arise (Fig. S1B). Fluorescent WISH and immunofluorescence analysis in *fli1a:GFP* embryos, which labels endothelial cells with GFP, showed that the expression of *jam3b* was partially overlapped with that of *fli1a:GFP* at 14 hpf (Fig. S1C), indicating that *jam3b* is expressed in endothelial precursors. The expression of *jam3b* was also detected in the DA, particularly in the ventral floor of the DA (Fig. S1D), the site of HSC emergence. These data indicate that *jam3b* is expressed in endothelial/hematopoietic tissues over the time frame of hemato-vascular development.

To investigate the role of Jam3b in the hemato-vascular development, we utilized a *jam3b* mutant zebrafish line, *jam3b^sa37^*, which contains a missense point mutation in a cysteine residue (C136Y) that is predicted to form a structurally critical disulphide bond (Powell et al., 2011). We first examined the expression of HSC marker genes, *runx1* and *cmyb*, in homozygous *jam3b^sa37^* embryos. The expression of *runx1* and *cmyb* was detected in the DA of wild type (WT) embryos, but expression of both genes was largely reduced in *jam3b^sa37^* embryos (Fig. 1A). Developing HSPCs can be visualized as *cd41:GFP*; *kdrl:mCherry* double-positive cells in the ventral floor of the DA (Clements et al., 2011). As shown in Fig. 1B, the number of double-positive cells was approximately four times lower in *jam3b^sa37^* embryos compared with WT embryos. After budding from the DA, nascent HSPCs migrate to and colonize the caudal hematopoietic tissue (CHT), an equivalent tissue to the mammalian fetal liver where HSPCs expand (Tamplin et al., 2015), and the thymus where T lymphocytes are produced (Trede and Zon, 1998). The expression of *cmyb* in the CHT and *rag1* (a T cell marker) in the thymus was nearly undetectable in *jam3b^sa37^* embryos at 4 days post-fertilization (dpf) (Fig. 1C). These results indicate that Jam3b is required for HSC formation in the developing embryo.

**Fig. 1.**
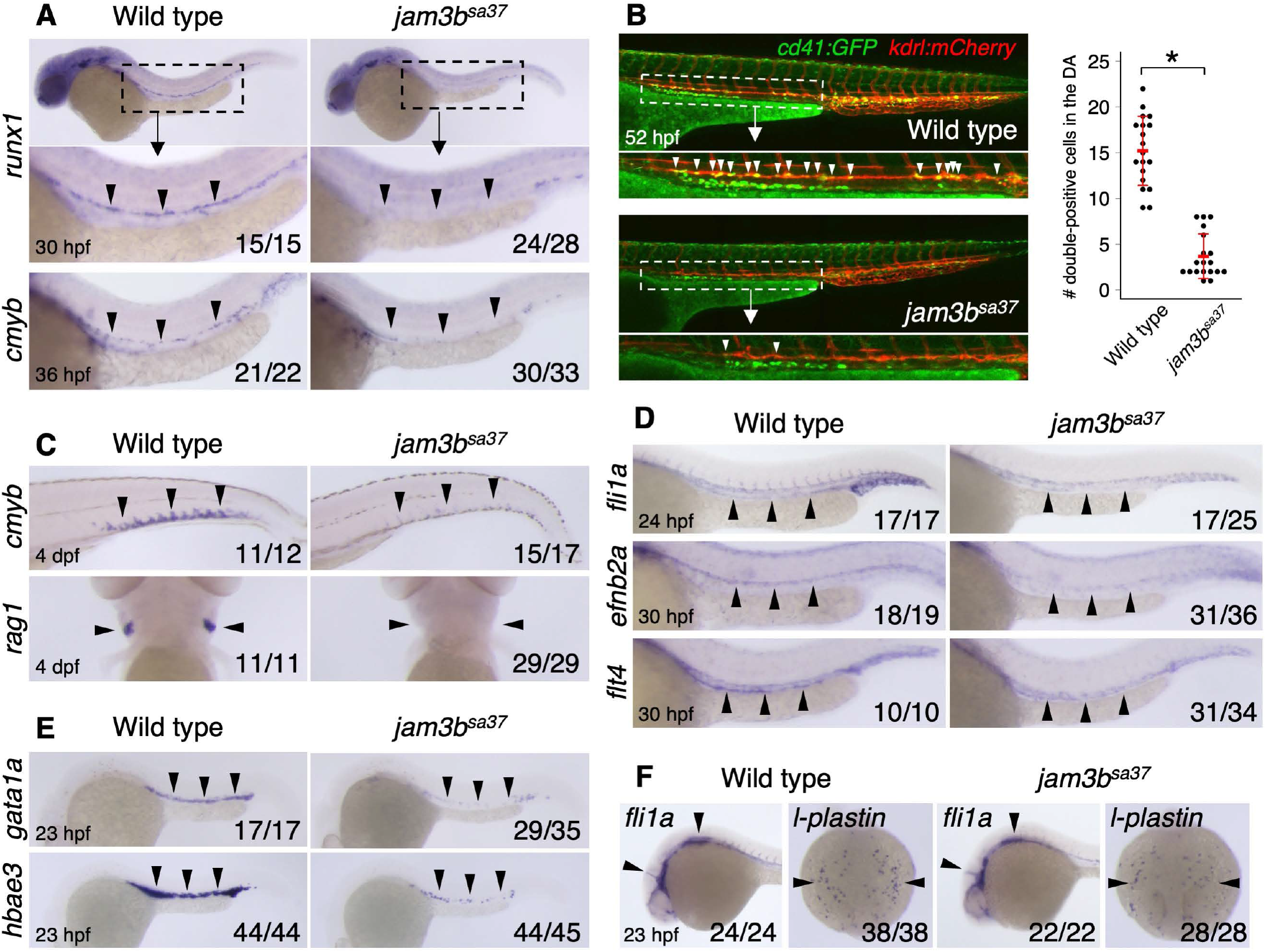
Mutation of *jam3b* causes defects in both hematopoietic and vascular development. (A) Expression of *runx1* and *cmyb* in the DA of WT or *jam3b^sa37^* embryos. (B) Number of *cd41:GFP*; *kdrl:mCherry* double-positive cells in WT or *jam3b^sa37^* embryos. White arrowheads indicate double-positive cells in the ventral floor of the DA. Mean ± s.d.. \**p* < 0.00001. (C) Expression of *cmyb* in the CHT and *rag1* in the thymus of WT or *jam3b^sa37^* embryos. (D) Expression of *fli1a* (endothelium), *efnb2a* (DA), and *flt4* (PCV) in WT or *jam3b^sa37^* embryos. (E) Expression of *gata1a* and *hbae3* in primitive erythrocytes of WT or *jam3b^sa37^* embryos. (F) Expression of *fli1a* and *l-plastin* (macrophage) in WT or *jam3b^sa37^* embryos. Black arrowheads indicate the expression domain of each gene (A, C-F).

In zebrafish, the N-terminal-truncated isoform of *scl*, *scl-β*, is expressed in hemogenic endothelial cells and shown to act as upstream of *runx1* (Qian et al., 2007; Zhen et al., 2013). A paralogue of zebrafish *gata2* genes, *gata2b*, is also known to regulate *runx1* expression in hemogenic endothelial cells (Butko et al., 2015). We found that the expression of *scl-β*, *gata2b*, and *gfi1aa*, an additional early HSC marker gene (Thambyrajah et al., 2016), was also remarkably reduced in the DA in *jam3b^sa37^* embryos (Fig. S1E), suggesting that mutation of Jam3b impairs the early specification program of HSCs.

We next investigated the formation of vascular endothelium and primitive hematopoietic cells in *jam3b^sa37^* embryos. In WT embryos, the expression of a pan-endothelial marker gene, *fli1a*, was detected in the whole vasculature including the DA, posterior cardinal vein (PCV), and intersegmental vessels (ISVs) at 24 hpf. In contrast, *fli1a* expression was weak and discontinuous in the trunk of *jam3b^sa37^* embryos at the same stage. We also examined the expression of *efnb2a* (a DA marker) and *flt4* (a PCV marker) in *jam3b^sa37^* embryos, and observed an expression pattern similar to *fli1a* in the DA and PCV, respectively (Fig. 1D). The expression of primitive erythrocyte marker genes, *gata1a* and *hbae3*, in the intermediate cell mass (ICM) and posterior blood island (PBI), which are the major sites of primitive erythropoiesis (Galloway and Zon 2003; Warga et al. 2009), was largely reduced in *jam3b^sa37^* embryos compared with WT embryos (Fig. 1E). Similar to *jam3b^sa37^* embryos, embryos injected with a *jam3b* morpholino oligo (MO), which blocks the translation of *jam3b* mRNA (Fig. S1F) (Powell et al., 2011), also showed a reduction in *runx1* expression in the DA, *fli1a* in the trunk vasculature, and *gata1a* in the ICM and PBI (Fig. S1G), confirming the results observed in *jam3b^sa37^* embryos. In contrast, the head vasculature and primitive macrophages were normally developed in *jam3b^sa37^* embryos as evidenced by the expression of *fli1a* in the head and *l-plastin* (a macrophage marker) in the rostral blood island (Fig. 1F), the major site of primitive myelopoiesis. Collectively, these data indicate that mutation of Jam3b results in a *cloche*-like phenotype, with defects in both hematopoietic and vascular development that are restricted to the posterior part of the embryo.

### Jam3b maintains the expression of *npas4l* in the PLPM

Given the defect of both hematopoietic and vascular development in *jam3b^sa37^* embryos, we speculated that Jam3b is involved in the early hemato-vascular development. We next examined, therefore, the expression of early hematopoietic/endothelial marker genes in *jam3b^sa37^* embryos. As shown in Fig. 2A, the expression of *gata1a*, *scl*, *etv2* (also known as *etsrp*), and *npas4l* was reduced in the PLPM in *jam3b^sa37^* embryos at 14 hpf compared with WT embryos. Quantitative polymerase chain reaction (qPCR) analysis in isolated *fli1a:GFP* (+) cells at 15 hpf confirmed the reduction of *npas4l* as well as *fli1a* expression in LPM cells of *jam3b^sa37^* embryos (Fig. 2B). We also observed the down-regulation of other LPM marker genes, *alas2*, *klf17*, *hhex*, and *ets1*, in *jam3b^sa37^* embryos, whereas expression of *cdx4* and its downstream target gene *lmo2* (Paik et al., 2013) was near normal or slightly up-regulated (Fig. S2A, B). In contrast to the PLPM, the expression of *spi1b* (a myeloid marker, also known as *pu.1*), *scl*, and *etv2* in the anterior LPM (ALPM), the site where primitive myeloid cells and endothelial precursors emerge, was not greatly affected in *jam3b^sa37^* embryos (Fig. 2C), which is consistent with our finding that Jam3b is dispensable for the formation of primitive myelocytes and head vasculature (Fig. 1F). The expression of *pax2a* (a pronephros marker) and *dlc* (a somite marker) was also unaffected in *jam3b^sa37^* embryos (Fig. S2B), suggesting that mutation of *jam3b* does not affect mesodermal induction.

**Fig. 2.**
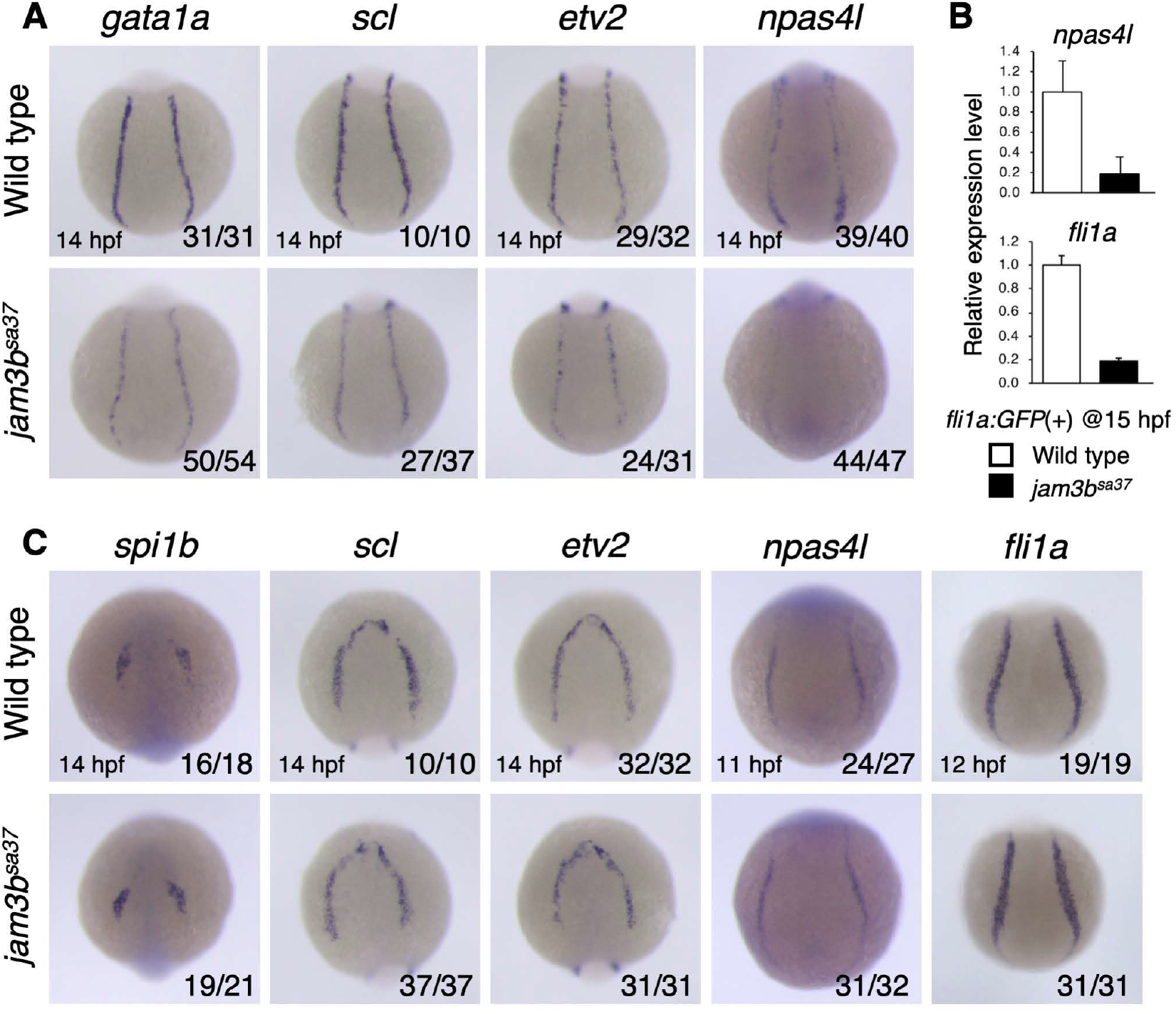
Mutation of *jam3b* reduces *npas4l* expression in the PLPM. (A) Expression of *gata1a*, *scl*, *etv2*, and *npas4l* in the PLPM of WT or *jam3b^sa37^* embryos. (B) Relative expression levels of *npas4l* and *fli1a* in isolated *fli1a:GFP* (+) cells from WT or *jam3b^sa37^* embryos. Error bars, s.d. (C) Expression of *spi1b*, *scl*, and *etv2* in the ALPM or *npas4l* and *fli1a* in the PLPM of WT or *jam3b^sa37^* embryos.

Since mutation of *jam3b* led to the reduction of various hematopoietic and endothelial marker genes including *npas4l* in the PLPM, we next questioned whether Jam3b is involved in the initial formation of the PLPM. The expression of *npas4l* in the PLPM is first detected at around 11 hpf as small bilateral stripes, which will extend anteriorly and posteriorly by 14 hpf (Reischauer et al., 2016). We found that *npas4l* expression was unaffected at 11 hpf in *jam3b^sa37^* embryos. Consistent with this, the expression of *fli1a* was also normal at 12 hpf (Fig. 2C). Taken together, these data suggest that Jam3b is required for the maintenance of *npas4l* expression in the PLPM rather than the PLPM formation.

### Jam3b regulates the expression of *drl* and *lrrc15*

In order to explore the molecular mechanisms governing Jam3b-dependent hemato-vascular development, RNA-seq analysis was conducted on WT and *jam3b^sa37^* embryos at 12 or 16 hpf (Fig. S3A). The principal component analysis (PCA) and correlation based on expression read counts revealed the close associations between each expression profile (Fig. S3B, C). Previous studies in zebrafish showed that *draculin* (*drl*), which encodes a zinc finger transcription factor, is expressed in the early LPM (Herbomel et al., 1999). Microarray analysis of isolated *drl:GFP* (+) and (-) cells at 11 hpf identified a number of uncharacterized genes expressed in the LPM (Mosimann et al., 2015). We therefore combined our RNA-seq datasets with the published microarray datasets of *drl:GFP* (+) and (-) cells to identify genes that were differentially expressed between LPM cells from WT and *jam3b^sa37^* embryos (Fig. S3A). Within genes that were over two-fold enriched in *drl:GFP* (+) cells (329 genes), 65 genes were selected as up-(> 2 fold) or down-regulated (< 0.5 fold) genes in either 12 or 16 hpf of *jam3b^sa37^* embryos. Among these 65 genes, 24 genes involved in signal transduction or encoding membrane proteins were further examined by qPCR in *fli1a:GFP* (+) cells at 15 hpf (Fig. S3D and Table S1). The expression of 16 out of the 24 selected genes was successfully detected by qPCR in a reproducible manner. The expression of *drl*, *gfi1b*, *lrrc15*, *mark4a*, *sox4a*, *sox32*, and *tcf21* was decreased in *fli1a:GFP* (+) cells of *jam3b^sa37^* embryos (< 0.3 fold), whereas the expression of *gsc* and *six1a* was increased (> 4 fold) (Fig. S3E). Among these 9 up- or down-regulated genes, we obtained *drl* and *lrrc15* as final candidates (Fig. 3A). *drl* was previously identified in a WISH screen of zebrafish cDNA libraries (Herbomel et al., 1999). There are four paralogues of *drl* genes in the zebrafish genome, *drl*, *drl-like 1* (*drl-l1*), *drl-l2*, and *drl-l3*, which contain three exons with very high homology in both coding and non-coding sequences. Although loss of *drl-l3*, which has the longest coding sequence of the group, affected primitive erythropoiesis, the role of *drl* as well as *drl-l1* and *drl-l2* in hemato-vascular development has not been clearly demonstrated (Pimtong et al., 2014). The expression of *drl* was specifically detected in the PLPM, but it was markedly decreased in *jam3b^sa37^* embryos at 15 hpf (Fig. 3B). Lrrc15 belongs to the leucine-rich repeat superfamily and is involved in cell-cell or cell-extracellular matrix interactions. Lrrc15 is frequently increased in tumor cells, particularly in breast and prostate cancers (Schuetz et al., 2006; Stanbrough et al., 2006). However, the role of Lrrc15 in hematopoietic or vascular cells has not been determined. In WT embryos, the expression of *lrrc15* was detectable by WISH after 17 hpf specifically in the vascular cord, a linear mass composed of endothelial precursors at the midline. The expression of *lrrc15* was also detected in the ventral floor of the DA and a portion of primitive erythrocytes at 24 hpf. We found that *lrrc15* expression was also largely decreased in the vascular cord at 18 hpf and the DA and primitive erythrocytes at 24 hpf in *jam3b^sa37^* embryos (Fig. 3C, D). Because the expression of *drl* and *lrrc15* was specific in the hematopoietic and/or endothelial tissue and was very strongly affected by mutation of *jam3b*, we hypothesized that Drl and Lrrc15 are core components of Jam3b-dependent hemato-vascular development.

**Fig. 3.**
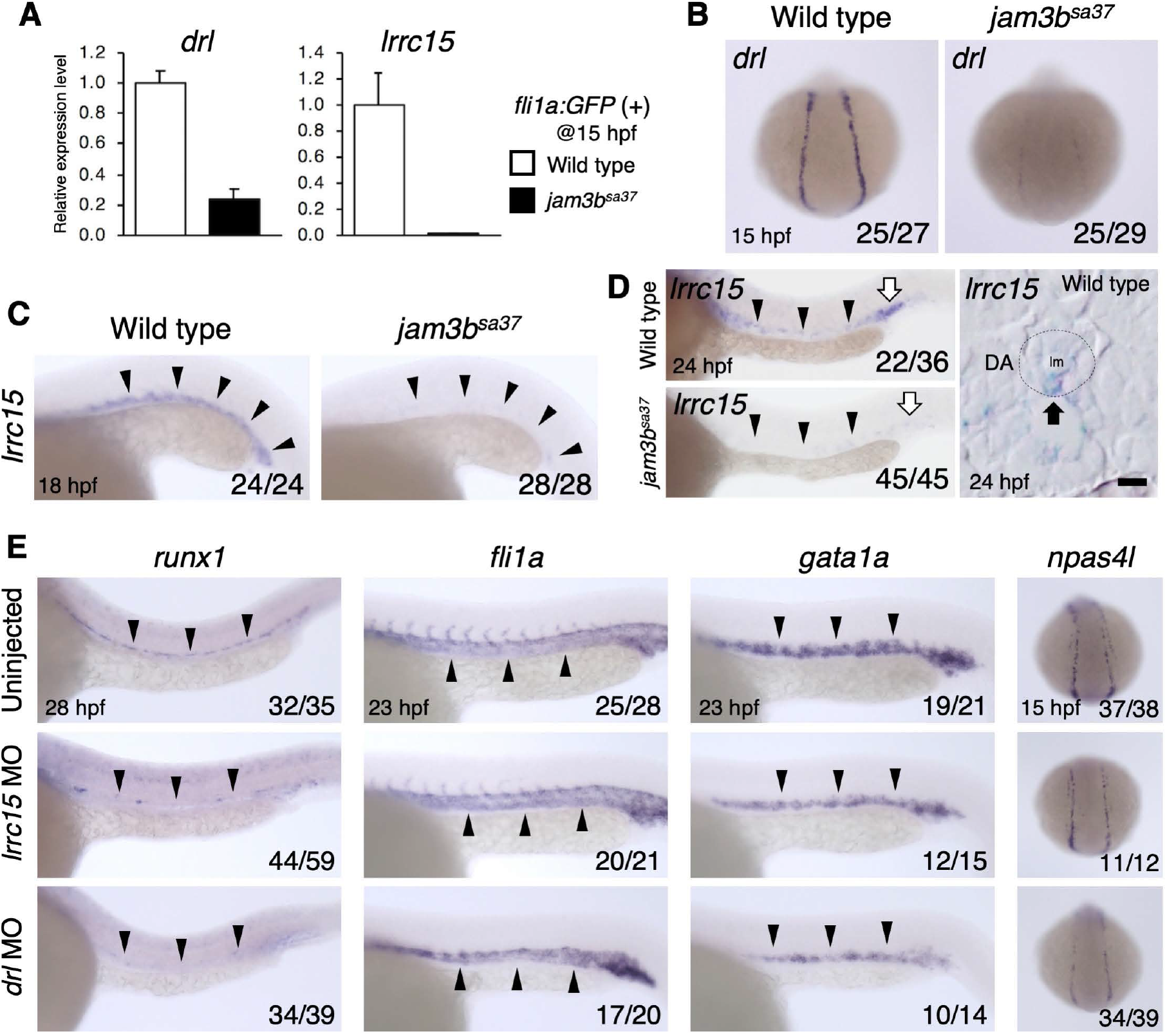
Transcriptome analysis identifies *drl* and *lrrc15* as a downstream target of Jam3b. (A) Relative expression levels of *drl* and *lrrc15* in isolated *fli1a:GFP* (+) cells from WT or *jam3b^sa37^* embryos. Error bars, s.d. (B) Expression of *drl* in the PLPM of WT or *jam3b^sa37^* embryos. (C, D) Expression of *lrrc15* in WT or *jam3b^sa37^* embryos. Black arrowheads indicate the vascular cord (C) or DA (D). White arrows indicate primitive erythrocytes in the PBI (D). A dotted circle outlines the DA, and a black arrow indicates an *lrrc15*-expressing cell in the ventral floor of the DA (D). lm, lumen; Bar, 5µm. (E) Expression of *runx1*, *fli1a*, *gata1a*, and *npas4l* in embryos uninjected or injected with *lrrc15* MO or *drl* MO. Black arrowheads indicate the expression domain of each gene.

To examine the functional roles of *drl* and *lrrc15* in hemato-vascular development, validated *drl* MO or *lrrc15* MO was injected into WT embryos (Fig. S4A, B) (Pimtong et al., 2014). In *lrrc15* morphants, the expression of *runx1* in the DA and *gata1a* in the ICM and PBI was reduced, whereas the expression of *fli1a* in the trunk vasculature and *npas4l* in the PLPM was unaffected. In contrast, the expression of all four of these genes was severely reduced in *drl* morphants (Fig. 3E). We generated *lrrc15* (*lrrc15^kz1^*) and *drl* (*drl^kz2^*) genetic mutant lines, which contain premature stop codons at early exons (Fig. S4C, D). Homozygous *lrrc15^kz1^* embryos exhibited reduced *runx1* and *gata1a* expression, but no change in *fli1a* expression. In contrast, the expression of all these three genes was reduced in homozygous *drl^kz2^* embryos (Fig. S4E), recapitulating both *lrrc15* and *drl* MO phenotypes. These results suggest that Lrrc15 is involved in hematopoietic development including formation of HSCs, whereas Drl is involved in both hematopoietic and vascular development.

To test whether the defects in hemato-vascular development in *jam3b^sa37^* embryos could be rescued by enforced expression of *drl* and/or *lrrc15*, we injected mRNAs for these two genes into *jam3b^sa37^* embryos. The expression of *npas4l* in the PLPM was increased in *jam3b^sa37^* embryos injected with *drl* mRNA compared with uninjected *jam3b^sa37^* embryos (Fig. S4F). Injection of *drl* mRNA also rescued the expression of *fli1a* in the trunk vasculature. However, the expression of *runx1* in the DA and *gata1a* in the ICM and PBI was not restored by injection of *drl* mRNA into *jam3b^sa37^* embryos (Fig. 4A), although *gata1a* expression increased in the PLPM (Fig. S4G). These data suggest that enforced expression of *drl* is sufficient to rescue vascular development, but insufficient to restore hematopoietic development in *jam3b^sa37^* embryos. In contrast to *drl* mRNA, injection of *lrrc15* mRNA rescued neither *runx1*, *fli1a*, nor *gata1a* in *jam3b^sa37^* embryos. Interestingly, however, co-injection of *drl* and *lrrc15* mRNA fully rescued the expression of *runx1* and *gata1a* as well as *fli1a* in *jam3b^sa37^* embryos (Fig. 4A). We examined the number of *cd41:GFP*; *kdrl:mCherry* double-positive HSPCs in the ventral floor of the DA and found that the average number of HSPCs in *jam3b^sa37^* embryos was significantly increased by co-injection of *drl* and *lrrc15* mRNA (Fig. 4B). In addition, the expression of *rag1* in the thymus was restored by co-injection of these two mRNAs (Fig. 4C). These data clearly indicate that enforced expression of both *drl* and *lrrc15* is sufficient to rescue the hemato-vascular development in *jam3b^sa37^* embryos. Our results also suggest that Drl acts as upstream of Npas4l to promote vascular development, while Lrrc15 is essential for hematopoietic development.

**Fig. 4.**
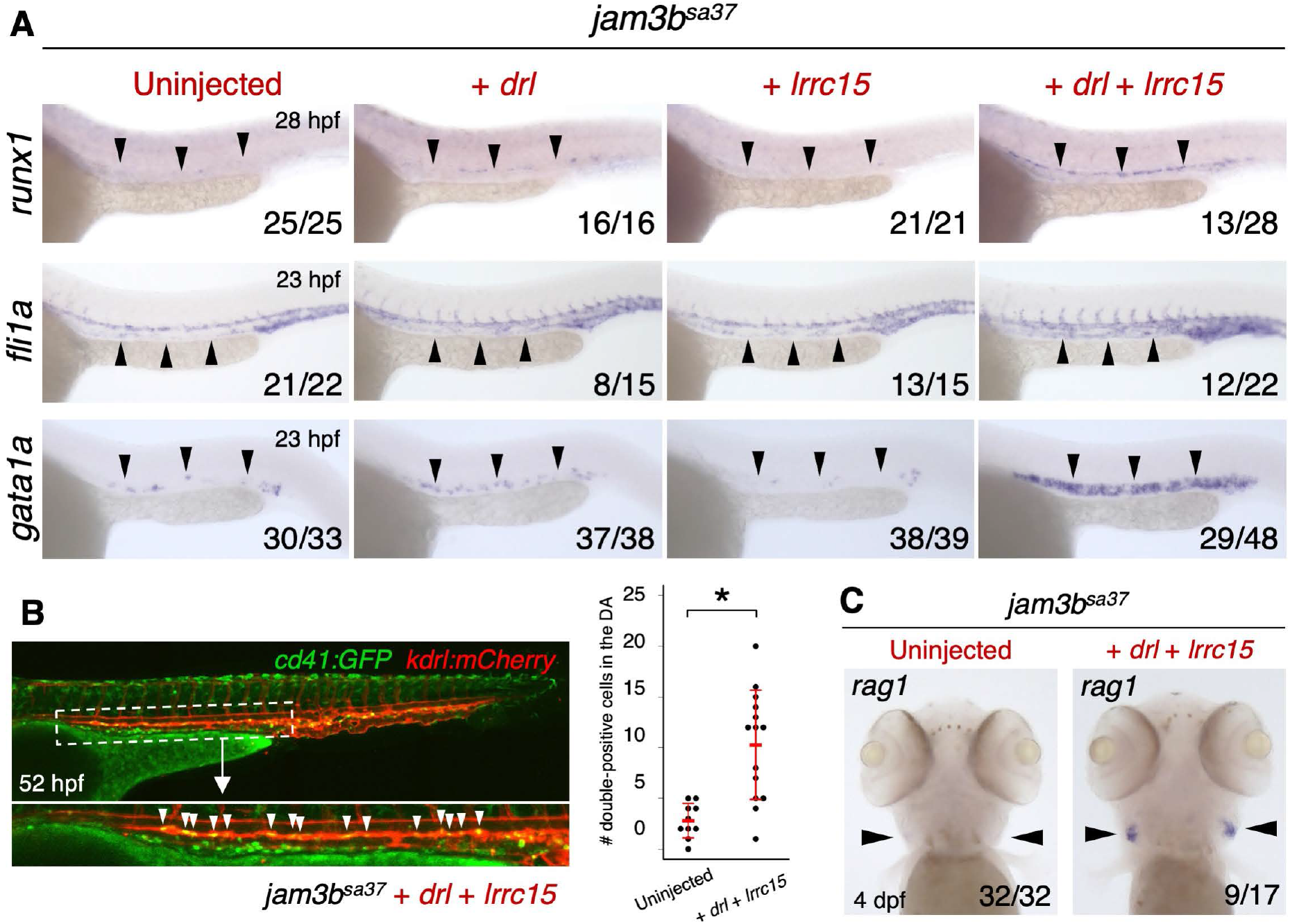
Co-injection of *drl* and *lrrc15* mRNA rescues hemato-vascular development in *jam3b^sa37^*embryos. (A) Expression of *runx1*, *fli1a*, and *gata1a* in *jam3b^sa37^* embryos uninjected or injected with *drl* mRNA, *lrrc15* mRNA, or both *drl* and *lrrc15* mRNA. (B) Number of *cd41:GFP*; *kdrl:mCherry* double-positive cells in *jam3b^sa37^* embryos co-injected with *drl* and *lrrc15* mRNA. White arrowheads indicate double-positive cells in the ventral floor of the DA. Mean ± s.d.. \**p* < 0.001. (C) Expression of *rag1* in the thymus in *jam3b^sa37^* embryos uninjected or co-injected with *drl* and *lrrc15* mRNA. Black arrowheads indicate the expression domain of each gene (A, C).

### Jam3b suppresses Rap1a-Erk activity to promote vascular development

In mammals, JAM3 has been shown to negatively regulate Rap1 activity (Orlova et al., 2006). Rap1 exists in an inactive guanine nucleotide diphosphate (GDP)-bound state and is activated when GDP is exchanged for guanine nucleotide triphosphate (GTP). Upon activation, Rap1 can bind to a variety of effectors, such as Raf family proteins, which triggers Erk activation and other signaling pathways (Stork and Dillon, 2005). We compared Rap1 activity between WT and *jam3b^sa37^* embryos by utilizing Raichu-Rap1, a fluorescence resonance energy transfer (FRET)-based Rap1 activation monitoring-probe. Within cells expressing Raichu-Rap1, binding of GTP-Rap1 to Raf induces FRET from CFP to YFP (Fig. S5A) (Sakurai et al., 2006). *Raichu-Rap1* mRNA was injected into WT or *jam3b^sa37^* embryos under the *kdrl:mCherry* background, which labels endothelial cells with mCherry, and live-imaging analysis was performed at 18 hpf. In WT embryos, FRET signals were nearly undetectable in either *kdrl:mCherry* (+) endothelial precursors and other cells. In contrast, strong FRET signals were detected in *jam3b^sa37^* embryos, especially in *kdrl:mCherry* (+) endothelial precursors within the posterior part of the embryo (Fig. 5A). This suggests that Jam3b suppresses Rap1 activation in the zebrafish embryo, as has been shown for human JAM3 (Orlova et al., 2006).

**Fig. 5.**
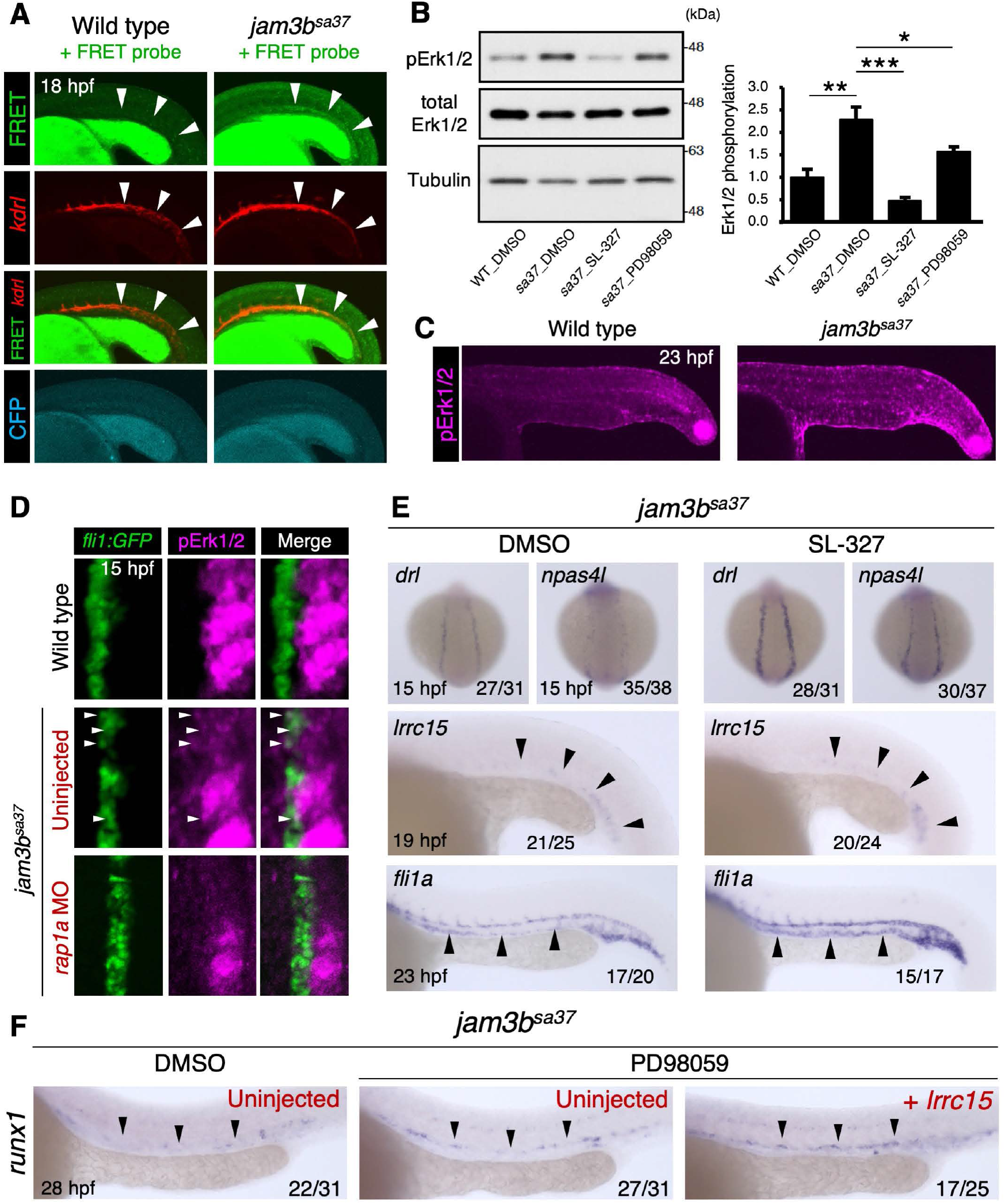
Jam3b suppresses Rap1a-Erk activity to promote hemato-vascular development. (A) Confocal imaging of FRET signals under the *kdrl:mCherry* background. WT and *jam3b^sa37^* embryos injected with *Raichu-Rap1* mRNA are shown. White arrowheads indicate the vascular cord labeled by *kdrl:mCherry*. (B) Western blotting analysis of phosphorylated Erk1/2 (pErk1/2), total Erk1/2, and Tubulin at 23 hpf in WT embryos treated with DMSO and *jam3b^sa37^* embryos treated with DMSO, SL-327, or PD98059. The graph denotes the ratio of pErk1/2 to total Erk1/2 in each embryo. Error bars, s.d.; \**p* < 0.05; \*\**p* < 0.01; \*\*\**p* < 0.001. (C) Immunostaining of pErk1/2 in WT or *jam3b^sa37^* embryos. The posterior part of embryos at 23 hpf are shown. (D) Immunostaining of pErk1/2 in WT or *jam3b^sa37^* embryos uninjected or co-injected with *rap1aa* MO and *rap1ab* MO. PLPM cells labeled with *fli1a:GFP* at 15 hpf are shown. White arrowheads indicate pErk1/2 (+) *fli1a:GFP* (+) cells. (E) Expression of *drl*, *npas4l*, *lrrc15*, and *fli1a* in *jam3b^sa37^* embryos treated with DMSO or SL-327. (F) Expression of *runx1* in the DA in *jam3b^sa37^* embryos treated with DMSO or PD98059 and uninjected or injected with *lrrc15* mRNA.

To determine if Jam3b regulates hemato-vascular development by suppressing Rap1 activity, Rap1 activation was induced by treating WT embryos with the Epac-selective cAMP analog 8-pCPT-2’-O-Me-cAMP-AM (known as “007-AM”). As shown in Fig. S5B, hyperactivation of Rap1 through 007-AM treatment led to the *cloche*-like phenotype observed in *jam3b^sa37^* embryos, marked by reduced expression of *drl*, *npas4l* in the PLPM, *runx1* in the DA, and *fli1a* in the trunk vasculature compared with DMSO-treated control embryos.

The Ras-Raf-Mek-Erk kinase cascade triggered by FGF signaling is one of the most important pathways for mesodermal induction in vertebrates (Umbhauer et al., 1995; Kimelman, 2006; Xiong et al., 2015). It has been shown that the duration and strength of Erk activation can influence cell fate determination, including proliferation, differentiation, survival, or apoptosis (Shaul and Seger, 2007). Because Rap1 is also known as a critical mediator of Erk activation in hematopoietic and vascular endothelial cells (Stork and Dillon, 2005; Yan et al., 2008), it is likely that the high level of Rap1 activation in *jam3b^sa37^* embryos causes excessive Erk activation, possibly leading to impaired hemato-vascular development. We therefore measured the level of phosphorylated Erk1 and 2 (pErk1/2), the active form of Erk1/2, in *jam3b^sa37^* embryos. We found that while the amount of total Erk1/2 was unchanged, pErk1/2 was higher in *jam3b^sa37^* embryos than WT embryos at 23 hpf. The higher ratio of pErk1/2 to total Erk1/2 in *jam3b^sa37^* embryos indicates that *jam3b* mutation results in increased Erk activation (Fig. 5B). Whole-mount immunostaining analysis further confirmed that pErk1/2 levels at 23 hpf were higher throughout the body of *jam3b^sa37^* embryos compared with WT embryos (Fig. 5C). At 15 hpf, pErk1/2 signals were strongly detected in the developing somite, but nearly undetectable in *fli1a:GFP* (+) PLPM cells in WT embryos, as has been shown previously (Shin et al., 2016). In contrast, pErk1/2 signals were detectable not only in the somite but also in a portion of *fli1a:GFP* (+) PLPM cells in *jam3b^sa37^* embryos (Fig. 5D). Importantly, MO knockdown of *rap1a* (both *rap1aa* and *rap1ab*) in *jam3b^sa37^* embryos reduced the level of Erk activation in PLPM cells (Fig. 5D, S5C), suggesting that Jam3b represses the Erk activation in PLPM cells through the regulation of Rap1a. MO knockdown of *rap1a* in WT embryos also led to the reduction of *drl* expression in the early mesoderm (Fig. S5D). These data suggest that Rap1a activates Erk to induce early mesodermal development, but this Rap1a-Erk signaling cascade is repressed later by Jam3b in PLPM cells.

To establish the link between Rap1a-Erk activation and the *cloche*-like phenotype in *jam3b^sa37^* embryos, we attempted to rescue the *jam3b^sa37^* phenotype by suppression of the Erk activity. We treated embryos with a chemical Mek inhibitor, either SL-327 or PD98059, to block Erk phosphorylation. Treatment of *jam3b^sa37^* embryos with 15µM of SL-327 or 10µM of PD98059 from 6 to 23 hpf reduced the phosphorylation of Erk1/2 (Fig. 5B). We observed a marked increase in *drl* and *npas4l* expression in the PLPM of SL-327-treated *jam3b^sa37^* embryos compared with DMSO-treated *jam3b^sa37^* embryos. Interestingly, however, the expression of *lrrc15* in the vascular cord was not restored by SL-327 treatment (Fig. 5E), indicating that the regulation of *lrrc15* by Jam3b is independent from the Erk signaling pathway. A previous study showed that the formation of ISVs, but not the DA or PCV, requires Erk activation in endothelial precursors (Shin et al., 2016). We found that the expression of *fli1a* was greatly increased in the DA and PCV in SL-327-treated *jam3b^sa37^* embryos, whereas it was unchanged in ISVs (Fig. 5E). Taken together, these data suggest that Jam3b represses the Rap1a-Erk signaling cascade to promote endothelial specification in mesodermal cells.

We next aimed to rescue HSC formation in *jam3b^sa37^* embryos by enforcing expression of *lrrc15* while blocking Erk signaling. It should be noted, however, that treatment with SL-327 severely affected the formation of the somite (Fig. S5E), which expresses a number of important signaling molecules that are required for HSC specification, such as Wnt16, Dlc, Dld, and Vegfa (Clements et al., 2011; Genthe and Clements, 2017). We found that the effect on somitogenesis was much less pronounced when embryos were treated with PD98059. Although PD98059 treatment could rescue the expression of *drl* and *npas4l* in the PLPM, it did not affect *lrrc15* expression in the vascular cord (Fig. S5E, F). We therefore injected *lrrc15* mRNA into *jam3b^sa37^* embryos and treated with PD98059 from 6 to 14 hpf. Consistent with our finding that *lrrc15* is required for HSC formation (Fig. 4A, S4E), the expression of *runx1* in the DA in *jam3b^sa37^* embryos was not recovered by PD98059 treatment. However, it was successfully rescued by injection of *lrrc15* mRNA together with PD98059 treatment (Fig. 5F). These data confirmed that Jam3b plays a second role in the acquisition of HSC fate through the regulation of *lrrc15* independently from *drl* regulation.

A previous study in mammals suggests that Rap1a-mediated Erk activation in vascular endothelial cells is induced at least in part by FGF signaling (Yan et al., 2008), raising the possibility that the high level of Erk activation observed in *jam3b^sa37^* embryos originates from FGF signaling. To test this hypothesis, we treated embryos with a chemical inhibitor of FGF receptors, SU5402. In WT embryos, treatment with 5µM of SU5402 from 8 to 14 hpf resulted in the enhanced expression of *drl* in the PLPM without causing morphological defects, whereas treatment from 2 to 14 hpf led to the deformation of the PLPM (Fig. S5G). Similar to Mek inhibitors, treatment of *jam3b^sa37^* embryos with SU5402 from 8 to 14 hpf rescued the expression of *drl* and *npas4l* in the PLPM and *fli1a* in the DA and PCV (Fig. S5H). Taken together, these results suggest that suppression of the FGF-Rap1a-Erk signaling pathway by Jam3b is crucial for the endothelial specification in mesodermal cells.

### Jam3b promotes HSC specification through an integrin-dependent signaling pathway

Having shown that Jam3b regulates *lrrc15* expression to promote HSC development independently from the Rap1a-Erk signaling cascade, we next questioned how Jam3b regulates *lrrc15* expression in endothelial precursors. In human carcinoma cells, JAM3 controls integrin-dependent signal transduction via the modulation of β1-integrin activation (Mandicourt et al., 2007). The activation of integrin via binding to extracellular matrix (ECM) mediates various intracellular signaling cascades, such as the phosphoinositide 3-kinase (PI3K), c-Jun N-terminal kinase (JNK), and Erk signaling cascade. Recently, it has been shown that β1-integrin is required for HSC specification in zebrafish embryos (Rho et al., 2019). To investigate if integrin regulates *lrrc15* expression in endothelial precursors, we utilized a transgenic zebrafish that expresses a dominant-negative form of β1-integrin fused with mCherry (mCherry-itgb1aDN). Enforced expression of *mCherry-itgb1aDN* leads to stacking of intracellular cytoskeleton-associated proteins, such as talin and actinin, which inhibits endogenous integrin signaling. (Iida et al., 2018). The *UAS:mCherry-itgb1aDN* line was crossed with an endothelial cell-specific Gal4 driver line, *fli1a:Gal4* (Fig. 6A). We found that the expression of *runx1* in the DA at 28 hpf and *lrrc15* in the vascular cord at 20 hpf was reduced in *mCherry-itgb1aDN* (+) embryos compared with *mCherry-itgb1aDN* (-) embryos (Fig. 6B, C). Although forced expression of *mCherry-itgb1aDN* in endothelial cells induced blood vessel abnormalities (Iida et al., 2018), the expression of *efnb2a* in the DA was intact in *mCherry-itgb1aDN* (+) embryos (Fig. S6A), suggesting that the defect of HSC formation in *mCherry-itgb1aDN* (+) embryos is not due to the secondary effect of blood vessel abnormalities.

**Fig. 6.**
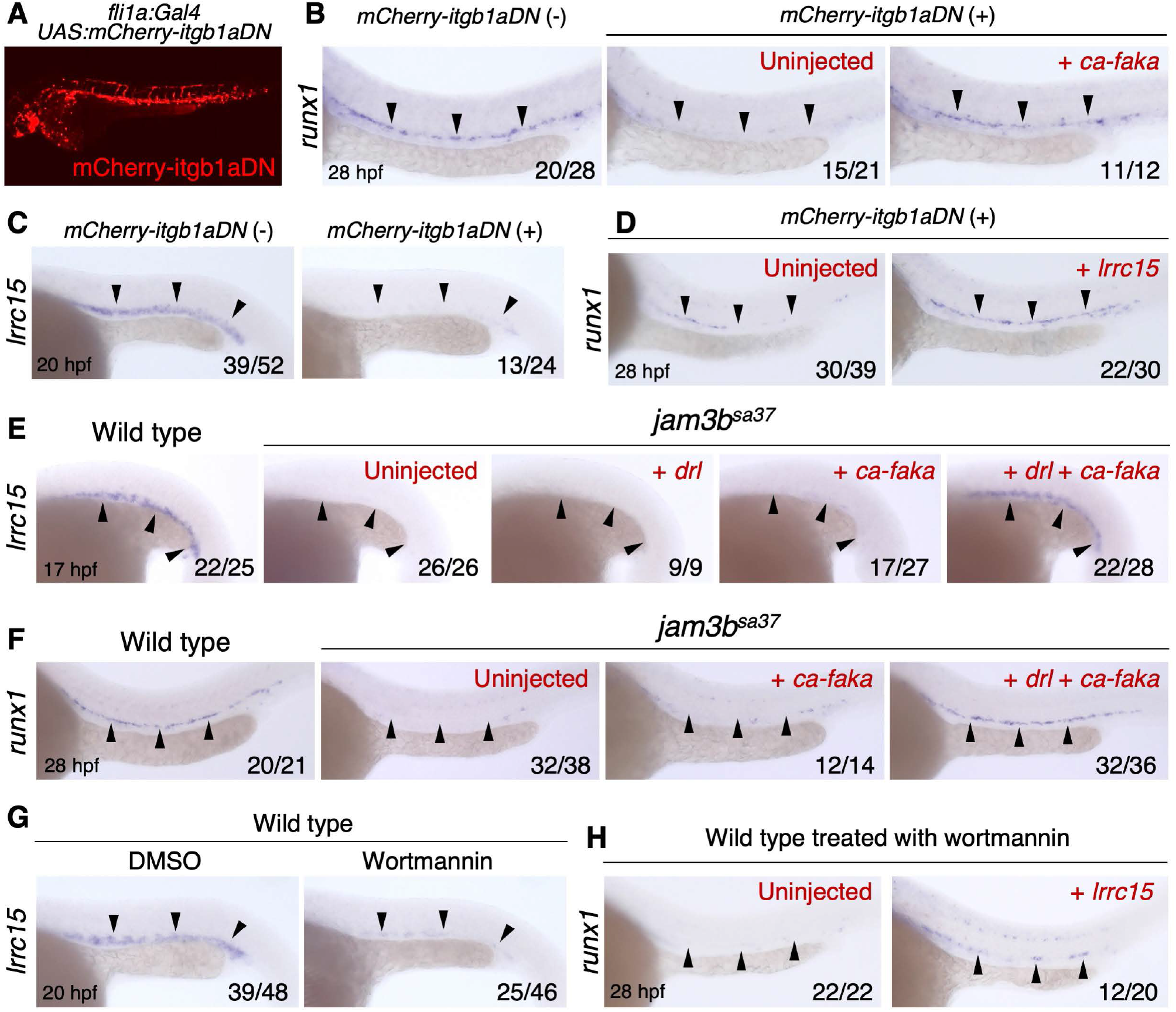
Jam3b regulates the integrin signaling pathway to form HSCs. (A) Fluorescent image of a *fli1a:Gal4*; *UAS:mCherry-itgb1aDN* embryo. (B) Expression of *runx1* in the DA of *mCherry-itgb1aDN* (-) or *mCherry-itgb1aDN* (+) embryos uninjected or injected with *ca-faka* mRNA. (C) Expression of *lrrc15* in the vascular cord of *mCherry-itgb1aDN* (-) or *mCherry-tgb1aDN* (+) embryos. (D) Expression of *runx1* in the DA of *mCherry-itgb1aDN* (+) embryos uninjected or injected with *lrrc15* mRNA. (E) Expression of *lrrc15* in the vascular cord of WT or *jam3b^sa37^* embryos uninjected or injected with *drl* mRNA, *ca-faka* mRNA, or both *drl* and *ca-faka* mRNA. (F) Expression of *runx1* in the DA of WT or *jam3b^sa37^* embryos uninjected or injected with *ca-faka* mRNA or both *drl* and *ca-faka* mRNA. (G) Expression of *lrrc15* in the vascular cord of WT embryos treated with DMSO or wortmannin. (H) Expression of *runx1* in the DA of WT embryos treated with wortmannin and uninjected or injected with *lrrc15* mRNA.

Focal adhesion kinase (FAK) is a nonreceptor tyrosine protein kinase that plays a central role in the transduction of integrin signaling. Upon binding of integrin to ECM, FAK is phosphorylated, which triggers recruitment of Src family kinases and signaling complex formation (Kleinschmidt and Schlaepfer, 2017). Mutation of lysine residues to glutamic acid in the activation loop of FAK (K578E/K581E) mimics phosphorylation of FAK and enhances integrin signaling (Gabarra-Niecko et al., 2002). We generated the mRNA for zebrafish constitutively activated Faka (*ca-faka*), which contains mutations of K580E and K583E (Fig. S6B). Injection of *ca-faka* mRNA was sufficient to rescue *runx1* expression in the DA in *mCherry-itgb1aDN* (+) embryos (Fig. 6B). In addition, injection of *lrrc15* mRNA also rescued *runx1* expression in *mCherry-itgb1aDN* (+) embryos (Fig. 6D). Together, these data suggest that integrin-dependent signaling regulates the expression of *lrrc15* in endothelial precursors to form HSCs in developing embryos.

To examine whether mutation of *jam3b* affects the activity of integrin in endothelial precursors, we isolated *fli1a:GFP* (+) cells from WT or *jam3b^sa37^* embryos at 18hpf and performed an adhesion assay on a fibronectin-coated dish. Although adherent *fli1a:GFP* (+) cells were detected in both WT and *jam3b^sa37^* embryos (Fig. S6C), the percentage of adherent *fli1a:GFP* (+) cells was significantly lower in *jam3b^sa37^* embryos compared with WT embryos (Fig. S6D), suggesting that mutation of *jam3b* down-regulates integrin-dependent interaction of endothelial precursors to ECM. These findings allowed us to test if HSC formation in *jam3b^sa37^* embryos can be rescued by injection of *ca-faka* mRNA. As shown in Fig. 6E, the expression of *lrrc15* in the vascular cord was restored in *jam3b^sa37^* embryos injected with both *drl* and *ca-faka* mRNA, whereas it was not restored in *jam3b^sa37^* embryos injected *ca-faka* mRNA alone. Consistent with this, the expression of *runx1* in the DA was also restored in *jam3b^sa37^* embryos injected with both mRNAs (Fig. 6F). These results strongly suggest that Jam3b regulates the integrin-dependent signaling cascade to promote HSC specification.

Since FAK is known to mediate multiple signaling pathways, including PI3K, JNK, and Erk, we then tried to determine which pathway is involved in HSC formation. We could exclude the Erk signaling pathway because impaired hemato-vascular development in *jam3b^sa37^* embryos is caused in part by the high level of Erk activation in PLPM cells (Fig. 5B-F). Therefore, we treated WT embryos with a PI3K inhibitor (wortmannin) or a JNK inhibitor (SP600125) from 6-24 hpf and examined *runx1* expression in the DA. We found that wortmannin treatment reduced *runx1* expression, while SP600125 treatment did not (Fig. S6E). Treatment of wortmannin also reduced *lrrc15* expression in the vascular cord, while enforced expression of *lrrc15* in wortmannin-treated embryos rescued *runx1* expression in the DA (Fig. 6G, H). Collectively, these data suggest that Jam3b plays a second role in the regulation of *lrrc15* to form HSCs via the activation of the integrin-FAK-PI3K signaling pathway.

Since heterophilic interactions between Jam2 and Jam3 have been implicated in many biological processes (Gliki et al., 2004; Weber et al., 2007; Powell et al., 2011; Arcangeli et al., 2011), we investigated whether Jam3b requires Jam2 to regulate hemato-vascular development. Previous studies determined the physical binding properties of zebrafish Jam proteins by surface plasmon resonance: Jam3b can bind to Jam2a, Jam2b, and Jam3b itself, but not Jam1a, Jam1b, and Jam3a (Powell et al., 2011; Powell et al., 2012). It has also been shown that the interaction of Jam1a and Jam2a mediates Notch signal transduction between endothelial precursors and the somite to establish HSC fate (Kobayashi et al., 2014), while the role of Jam2b in hemato-vascular development is not determined. MO knockdown of *jam2a*, but not *jam2b*, reduced the expression of *drl* in the PLPM and *lrrc15* in the vascular cord (Fig. S7A, B). Although the phenotype of *jam2a* morphants was similar to that of *jam3b^sa37^* embryos, *jam2a* expression was not detected in the PLPM (Fig. S7C), suggesting that non-endothelial Jam2a is involved in the activation of Jam3b in the PLPM to promote hemato-vascular development. In contrast to *jam3b^sa37^* embryos, however, *runx1* expression in the DA was not restored in embryos injected with *jam2a* MO together with *drl* and *lrrc15* mRNA (Fig. S7D). This is likely due to the additional role that Jam2a plays in regulating HSC development via Notch signal transduction (Kobayashi et al., 2014). Taken together, our data suggest that Jam3b regulates both endothelial and hematopoietic specification through two independent signaling pathways in a Jam2a-dependent manner (Fig. 7).

**Fig. 7.**
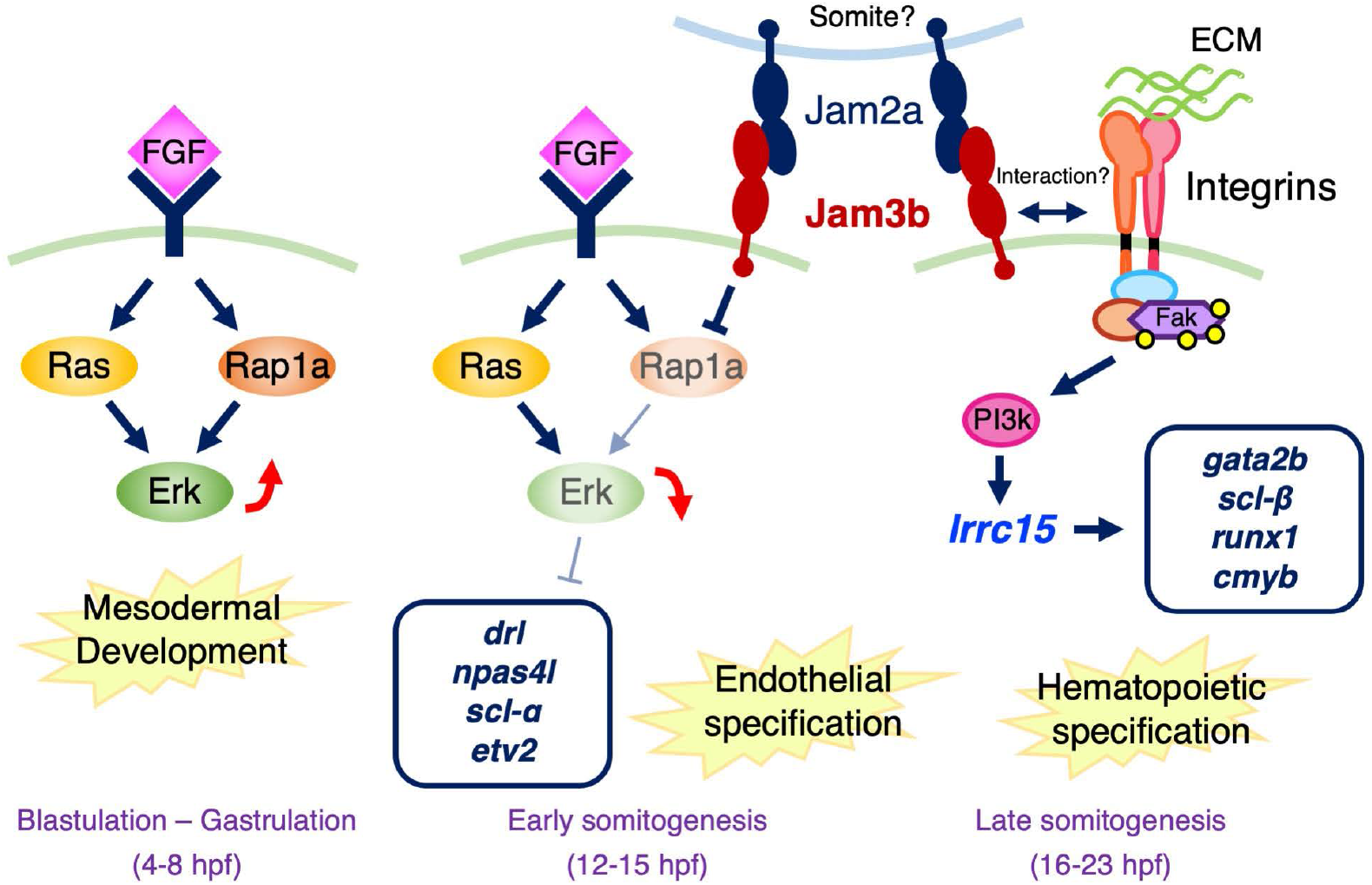
Hemato-vascular development in the zebrafish embryo. Schematic diagram of hemato-vascular development in the zebrafish embryo. During blastulation to gastrulation, FGF signaling induces mesodermal development via the activation of Erk. This Erk activation in mesodermal cells is suppressed by Jam3b through inhibition of the Rap1a activity, leading to the expression of endothelial-related genes, such as *drl*, *npas4l*, *scl-α*, and *etv2*. Jam3b then regulates the expression of *lrrc15* in endothelial precursors via an integrin-dependent signaling pathway to drive hematopoietic-related genes, such as *gata2b*, *scl-β*, *runx1*, and *cmyb*. The regulation of hemato-vascular development by Jam3b is dependent on Jam2a expression

## Discussion

In the present study, we define a novel molecular mechanism by which Jam3b plays dual roles in both hematopoietic and vascular development in mesodermal cells. During early embryogenesis, Erk is activated by FGF signaling to induce mesodermal development. This default state of Erk activation in mesodermal cells is suppressed by Jam3b through inhibition of the Rap1a activity, promoting the acquisition of early endothelial identity. Jam3b then regulates the expression of *lrrc15* in endothelial precursors via an integrin-dependent signaling pathway to confer hematopoietic fate, leading to the formation of HSCs in the DA. Thus, our study delineates the fundamental mechanisms for the divergence of hematopoietic and vascular lineages from mesodermal cells.

Jam proteins play major roles in the modulation of cell-cell interaction, cell migration, and cell polarization in many tissues. In some cases, however, Jam proteins are also involved in the signal transduction to promote cell proliferation and differentiation. For instance, Jam1 forms a complex with integrin α_V_β_3_ in endothelial cells under physiological conditions. After activation of endothelial cells by FGF signaling, Jam1 dissociates from integrin α_V_β_3_, which induces angiogenetic signals, such as the Erk signaling pathway (Naik et al., 2003). Our current data and a previous study in mammals (Orlova et al., 2006; Mandicourt et al., 2007) demonstrated that Jam3 negatively regulates the Rap1a signaling pathway, but positively regulates integrin-dependent signaling pathway. Although further studies are required to address the detailed molecular mechanism behind the regulation of Rap1a and integrin by Jam3, these regulatory pathways may be evolutionarily conserved among vertebrates.

Identification of *npas4l* has greatly contributed to our understanding of how hematopoietic/endothelial precursors are specified from the mesodermal cell in vertebrates (Reischauer et al., 2016). In the present study, we have demonstrated that Drl acts as upstream of *npas4l* in the LPM. Loss of *drl* results in the down-regulation of *npas4l*, whereas enforced expression of *drl* increased *npas4l* expression in the PLPM of *jam3b^sa37^* embryos. Interestingly, injection of *drl* mRNA did not lead to the ectopic expression of *npas4l* or other hematopoietic/endothelial marker genes, possibly due to the presence of a cofactor that regulates *npas4l* expression in the LPM. On the other hand, *drl* expression is decreased in the PLPM by the loss of *scl* or *lmo2* (Dooley et al., 2005; Patterson et al., 2007), two downstream targets of Npas4l (Reischauer et al., 2016). These findings, in combination with our data, suggest that there are cross-talk regulatory mechanisms between Drl, Scl, Lmo2, and Npas4l in the PLPM, and these factors may cooperatively regulate early hemato-vascular development.

The duration and strength of Erk activation is tightly controlled by the balance between positive and negative regulatory pathways. Imbalanced Erk signaling is frequently associated with malignant transformation. Megakaryocyte maturation is regulated by sustained Erk activation via Rap1 activation, whereas it is inhibited by transient Erk activation via Ras activation (Stork and Dillon, 2005). These observations highlight that sustained or transient activation of Erk signaling can direct different cellular outcomes. In the present study, we found that mutation of *jam3b* results in the hyperactivation of the Rap1a-Erk signaling cascade in PLPM cells. While Erk activation induced by FGF signaling is required for early mesodermal induction, prolonged Erk activation in the PLPM impaired hemato-vascular development. Thus, our data suggest that Jam3b plays an important role in the repression of Rap1a activity, enabling the transient Erk activation in the PLPM to promote endothelial specification.

In vertebrates, it is believed that hemangioblasts can give rise to hematopoietic cells through two types of intermediate endothelial precursors, hemogenic angioblasts and hemogenic endothelial cells. Hemogenic angioblasts can give rise to primitive hematopoietic cells in the yolk sac, and hemogenic endothelial cells can generate HSCs in the DA (Gritz and Hirschi, 2016; Lacaud and Kouskoff, 2017). This concept suggests that the endothelial specification is an essential first step for mesodermal cells to differentiate into hematopoietic lineages. Consistent with this idea, our present data suggest that after promoting endothelial specification through Npas4l regulation, Jam3b plays a second role in the acquisition of hematopoietic fate by regulating *lrrc15*. Although the role of Lrrc15 in HSC formation remains elusive, our work uncovers the importance of Jam3 in the acquisition of both endothelial and hematopoietic fates by mesodermal cells, opening new avenues for the generation of HSCs from iPSCs.

## Materials and methods

### Zebrafish husbandry

Zebrafish strains, AB*, *jam3b^sa37^*, *lrrc15^kz1^*, *drl^kz2^*, *Tg(fli1a:GFP)^y1^*, *Tg(−6.0itga2b:eGFP)^la2^*, *Tg(kdrl:memCherry)^s896^*, *Tg(fli1a:Gal4FF)^ubs4^*, *Tg(UAS:mCherry-itgb1aDN)^ko112^* were raised in a circulating aquarium system (AQUA) at 28.5°C in a 14/10 h light/dark cycle and maintained in accordance with guidelines of the Committee on Animal Experimentation of Kanazawa University. Because homozygous *jam3b^sa37^* animals are partially viable into adulthood and fertile, homozygous *jam3b^sa37^* embryos were obtained by an in-cross of *jam3b^sa37^* animals and were used for experiments.

### Chemical treatment

All chemical compounds used in this study were dissolved in dimethyl sulfoxide (DMSO) (Wako) and were used at the following concentration: 8-pCPT-2’-O-Me-cAMP-AM (007-AM) (Biolog Life Science Institute), 10 µM; SU5402 (Wako), 5 µM; SL-327 (Santa Cruz), 15 µM; PD98059 (Tocris Bioscience), 10 µM; wortmannin (Tocris Bioscience), 1 µM; SP600125 (Tocris Bioscience), 5 µM. Embryos were incubated in E3 medium (5 mM NaCl, 0.17 mM KCl, 0.33 mM CaCl_2_, 0.33 mM MgSO_4_) containing DMSO or a chemical compound in the dark, followed by dechorionation with 20 mg/mL of pronase (Sigma) in E3 medium and fixation with 4% paraformaldehyde (PFA) (Wako) in phosphate buffered saline (PBS).

### Generation of mutant lines

For CRISPR/Cas9-mediated generation of mutant lines, The crRNA recognizing the genomic loci was designed and purchased from Integrated DNA Technologies (IDT) and hybridized with tracrRNA according to the procedures from IDT. The gRNA complex (50 pg/nL) was then co-injected with 400 ng/nL of Cas9 protein (IDT) into one-cell stage embryos. Fish from the F1 generation were screened using primers that flanked the target region (Table S2), and mutant alleles were identified by sequencing. Embryos from the F3 generation were used for experiments. The crRNA sequences used in this study are as follows: *lrrc15*, CTGGCAGCTTTCTGGACATG; *drl*, ACAGAGCATCACAATGCTGA.

### Whole-mount *in situ* hybridization and immunostaining

cDNAs were cloned by reverse transcription (RT)-PCR using specific primers listed in Table S2, and ligated into the pCRII-TOPO vector (Invitrogen). Digoxigenin (DIG)-labeled RNA probes were prepared by *in vitro* transcription with linearized constructs using DIG RNA Labeling Kit (SP6/T7) (Sigma). Embryos were fixed in 4% PFA in PBS. For permeabilization, embryos were treated with proteinase K (10 µg/ml) (Sigma) for 30 sec to 30 min, refixed with 4% PFA, and washed with 0.1% Tween-20 (Sigma) in PBS (PBST). Hybridization was then performed using DIG-labeled antisense RNA probes diluted in hybridization buffer (50% formamide, 5X standard saline citrate (SSC), 0.1% Tween-20, 500 µg/mL torula RNA, 50 µg/mL heparin) for 3 days at 65°C. For detection of DIG-labeled RNA probes, embryos were blocked in 0.2% bovine serum albumin (BSA) (Sigma) in PBST and incubated overnight at 4°C with the alkaline phosphatase-conjugated anti-DIG antibody (Sigma) at a dilution of 1:5000. After washing with PBST, embryos were developed using nitroblue tetrazolium chloride and 5-bromo-4-chloro-3-indolyl-phosphate (NBT/BCIP) (Sigma) in staining buffer (100 mM Tris pH9.5, 100 mM NaCl, 50 mM MgCl_2_, 0.1% Tween-20). For histological analysis, embryos developed with NBT/BCIP were embedded in paraffin and sectioned at 3 µm in thickness. Deparaffinized tissue sections were mounted with Mount-Quick Aqueous (Cosmo Bio). For fluorescent whole-mount *in situ* hybridization, embryos were permeabilized with acetone at −20°C for 7 min and hybridized with DIG-labeled RNA probes as described above. Embryos were blocked with 2% blocking reagent (Sigma) in PBST and incubated with the peroxidase-conjugated anti-DIG antibody (Sigma) at a dilution of 1:500. Embryos were developed using TSA Plus Cyanine 3 System (Perkin Elmer) for 30 min. For whole-mount immunostaining, fixed embryos were permeabilized with acetone, blocked with 2% blocking reagent, and incubated overnight at 4°C with the rabbit anti-phospho-p44/42 MAPK (for pErk1/2) (Cell Signaling Technology) at 1:250, and/or chicken anti-GFP (Aves) at 1:1000 dilution. After washing with PBST, embryos were incubated overnight at 4°C with the goat anti-chicken IgY Alexa Fluor 488-conjugated (Abcam) at 1:1000 and/or donkey anti-rabbit IgG Alexa Fluor 647-conjugated (Abcam) at 1:1000 dilution.

### Cell preparation and flow cytometry

Embryos were dechorionized and digested with 50 µg/mL of Liberase TM (Roche) in PBS for 1 hour at 37°C. Cells were then filtered through a 40 µm stainless steel mesh and washed with 2% fetal bovine serum (FBS) (Cosmo Bio) in PBS by centrifugation. Just before flow cytometric analysis, Sytox Red (Thermo Fisher) solution was added to the samples at a final concentration of 5 nM to exclude nonviable cells. Flow cytometric acquisition and cell sorting was performed on a FACS Aria II (BD Biosciences). Cells were sorted into 10% FBS in PBS and were used for expression analysis.

### Adhesion assay

Sorted *fli1a:GFP* (+) cells were resuspended in the medium containing 40% Leiboviz’s L-15 medium (Wako), 32% Dulbecco’s modified Eagle’s medium (Wako), 12% Ham’s F12 medium (Wako), 8% FBS, 2mM L-glutamine (Wako), 15mM 4-(2-hydroxyethyl)-1-piperazineethanesulfonic acid (HEPES, Sigma), 100U penicillin (Wako), and 100 µg/mL streptomycin (Wako). Cells were then plated on a 96- well plate (Violamo) or a glass-bottom dish (Matsunami) coated with 50 µg/mL of fibronectin (Cosmo Bio) and incubated for 12 hrs at 30°C, 5% CO_2_. For the adhesion assay, cells in a 96-well plate were washed with PBS three times and GFP (+) adherent cells were counted using an EVOS fl (Thermo Fisher Scientific). Percent adhesion was calculated by dividing the number of GFP (+) adherent cells by the total number of GFP (+) cells plated in the well.

### qPCR and RT-PCR

Total RNA was extracted from embryos using RNeasy Mini Kit (Qiagen), and cDNA was synthesized using ReverTra Ace qPCR RT Master Mix (Toyobo). Quantitative real-time PCR (qPCR) assays were performed using GoTaq qPCR Master Mix (Promega) on a ViiA 7 Real-Time PCR System according to manufacturer’s instructions (Thermo Fisher Scientific). The expression of *ef1a* was used for normalization. RT-PCR was performed using GoTaq Green Master Mix (Promega), and gel analysis was performed in 1.5% agarose or 5 - 10% acrylamide gels. Primers used for qPCR or RT-PCR are listed in Table S2.

### RNA-seq

Cap analysis of Gene Expression (CAGE) library preparation, sequencing, mapping, and gene expression analysis were performed by Dnaform (Haberle et al., 2015). Two micrograms of total RNA from embryos were used for synthesis of cDNA using a Library Preparation kit (Dnaform). 5’ caps of RNAs were biotinylated after reverse transcription. RNA-cDNA hybirds were then captured by streptavidin-conjugated magnetic beads. The cDNA was released from RNAs and ligated with 5′ and 3’ linkers. Second-strand synthesis was performed to create the final double-stranded DNA product. Sequencing of CAGE tags was performed using the Illumina NextSeq500, and base-calling was performed using the Illumina RTA software (ver. 2.4.11). Sequenced CAGE tags were mapped to the full genome sequences for *Danio rerio* (danRer10) using the Burrows-Wheeler Aligner (BWA, ver. 0.7.12-r1039) and HiSAT2 software (ver. 2.0.5). Tags per million (TPM) were calculated using the CAGEr package of Bioconductor in R ver. 3.3.3 with a cutoff of < 0.5. To evaluate quality and reproducibility, Pearson correlation coefficients were calculated and principal components analysis was performed using R. Hierarchical clustering of each subset was performed in R with the Bioconductor pvclust package. The data have been deposited in Gene Expression Omnibus (GEO) (National Center for Biotechnology Information) and are accessible through the GEO database (series accession number, GSE114416). Microarray data of *drl:GFP* (+) and *drl:GFP*(-) were obtained from the GEO database (series accession number, GSE70881).

### Western blotting

Dechorionized embryos were collected and lysed in lysis buffer (25 mM Tris-HCl pH7.4, 1 mM EDTA, 0.1 mM EGTA, 150 mM NaCl, 5 mM MgCl_2_, 2 mM Na_3_VO_4_, 20% glycerol, 0.1% Triton X-100, 1 mM dithiothreitol, proteinase inhibitor). The lysate was then centrifuged to remove debris, resuspended in 2X sample buffer (4% SDS, 0.2 M dithiothreitol, 0.1 M Tris-HCl pH6.8, 10% glycerol, 20 mg/mL bromophenol blue), and boiled to prepare SDS-PAGE samples. The samples were separated by a 10% polyacrylamide gel and transferred to a PVDF membrane (Millipore). The membrane was blocked with the Can Get Signal / PVDF blocking reagent (Toyobo) for 60 min at 37°C, and then incubated with rabbit anti-phospho-p44/42 MAPK (for pErk1/2; 1:500), rabbit anti-p44/42 MAPK (for total Erk1/2; Cell Signaling Technology; 1:1000), or rabbit anti-α/β Tubulin (Cell Signaling Technology; 1:1000) antibodies overnight at 4°C. After washing with TBS-T (50 mM Tris-HCl pH7.4, 138 mM NaCl, 2.7 mM KCl, 0.1% Tween-20), the membrane was incubated with the anti-rabbit IgG HRP-conjugated secondary antibody (Cosmo Bio; 1:10000) for 2 hours at room temperature. The primary and secondary antibodies were diluted with the Can Get Signal / Solution-1 and −2 (Toyobo), respectively. After washing with TBS-T, chemiluminescence reaction was performed using the Western Lightning Plus-ECL (Perkinelmer), and the signals were developed on X-ray films. Expression levels were quantified by the ImageJ software (ver. 1.51s).

### Microscopy

For fluorescent imaging, embryos were mounted in a glass bottom dish filled with 0.6% low-gelling agarose (Sigma) in E3 medium containing 200 µg/mL of tricaine (Sigma) and imaged using an FV10i confocal microscope and Fluoview FV10i-SW software (ver. 2.1.1) (Olympus). Visible light imaging or fluorescent images of GFP mRNA-injected embryos were captured using an Axiozoom V16 microscope (Zeiss) with a TrueChrome II digital camera (BioTools) and TCapture software (ver. 4.3.0.602) (Tucsen Photonics) or an Axioskop 2 plus microscope (Zeiss) with a DS74 digital camera and CellSens Standard software (ver. 1.16) (Olympus).

### Morpholino and mRNA injection

Embryos were injected at the one-cell stage with 1 nl of morpholino oligonucleotides (MO, GeneTools) or mRNA. The MO sequences and concentrations used in this study are as follows: *jam3b* MO, TTAACGCCATCTTGGAGTCGGTGAA (3.5 ng/nL) (Powell et al., 2011); *lrrc15* MO, AACCACGCCAGGTCCATTGATTCCG (1.5 ng/nL); *drl* MO, TCAAGAACAGACACCAACCTCTTTG (3 ng/ nL) (Pimtong et al., 2014); *rap1aa* MO, TGGTGGCCTGTAAGATTACATCACA (1 ng/nL); *rap1ab* MO, GTACAATGTACTTACCAAGGCTGAC (4 ng/nL). *jam2a* MO, AGGAACTACAGCAGAAACAGGTCAA (3.5 ng/nL) (Kobayashi et al., 2014); *jam2b* MO, GGTTAAAGCAATGCACCAACCAGAC (13 ng/nL). The *Raichu-Rap1* was amplified using *pCAGGS-Raichu-Rap1* vector, which consisted of *EYFP-V68L/Q69K*, zebrafish *rap1b*, *Raf RBD* and *ECFP* from the amino terminus, and was ligated into the *pCS2+* vector. Capped mRNAs were synthesized from linearized *pCS2+* constructs using the mMessage mMachine SP6 kit (Thermo Fisher Scientific). mRNAs were injected into embryos at following concentrations; *drl*, 50 pg/nL; *lrrc15*, 280 pg/nL; *Raichu-Rap1*, 50 pg/nL; *jam3b-GFP*, 50 pg/nL; *lrrc15-GFP*, 50 pg/nL; *ca-faka*, 100 pg/nL.

### Quantification and statistical analyses

Data were analyzed for statistical significance after at least two repeated experiments. Statistical differences between groups were determined by unpaired two-tailed Student’s *t*-test. A value of *p* < 0.05 was considered to be statistically significant.

## Acknowledgements

The authors thank G. Wright for providing the *jam3b^sa37^* line, D. Traver for providing the *fli1a:GFP*, *cd41:GFP*, and *kdrl:mCherry* lines, A. Iida for providing the *UAS:mCherry-itgb1aDN* line, M. Kobayashi for providing *in situ* probes, D. Voon and M. Hazawa for supporting experiments, and D. Voon and K. Lewis for critical evaluation of the manuscript.

## Author contributions

I.K. designed the research and experiments, performed flow cytometric analyses, histological analyses, imaging analyses, and bioinformatics analyses, and wrote the manuscript. I.K., J.K-S., Y.H. M.O. and K.Y. performed microinjection, expression analyses, generated mutant lines, and genotyped animals. H.K. performed western blotting, S.F. designed and generated the Raichu-Rap1 probe. I.K., H.K., S.F., and M.Y. discussed results and analyzed data, and H.K. and M.Y. edited the manuscript.

## Supplemental Figure legends

**Fig. S1.**
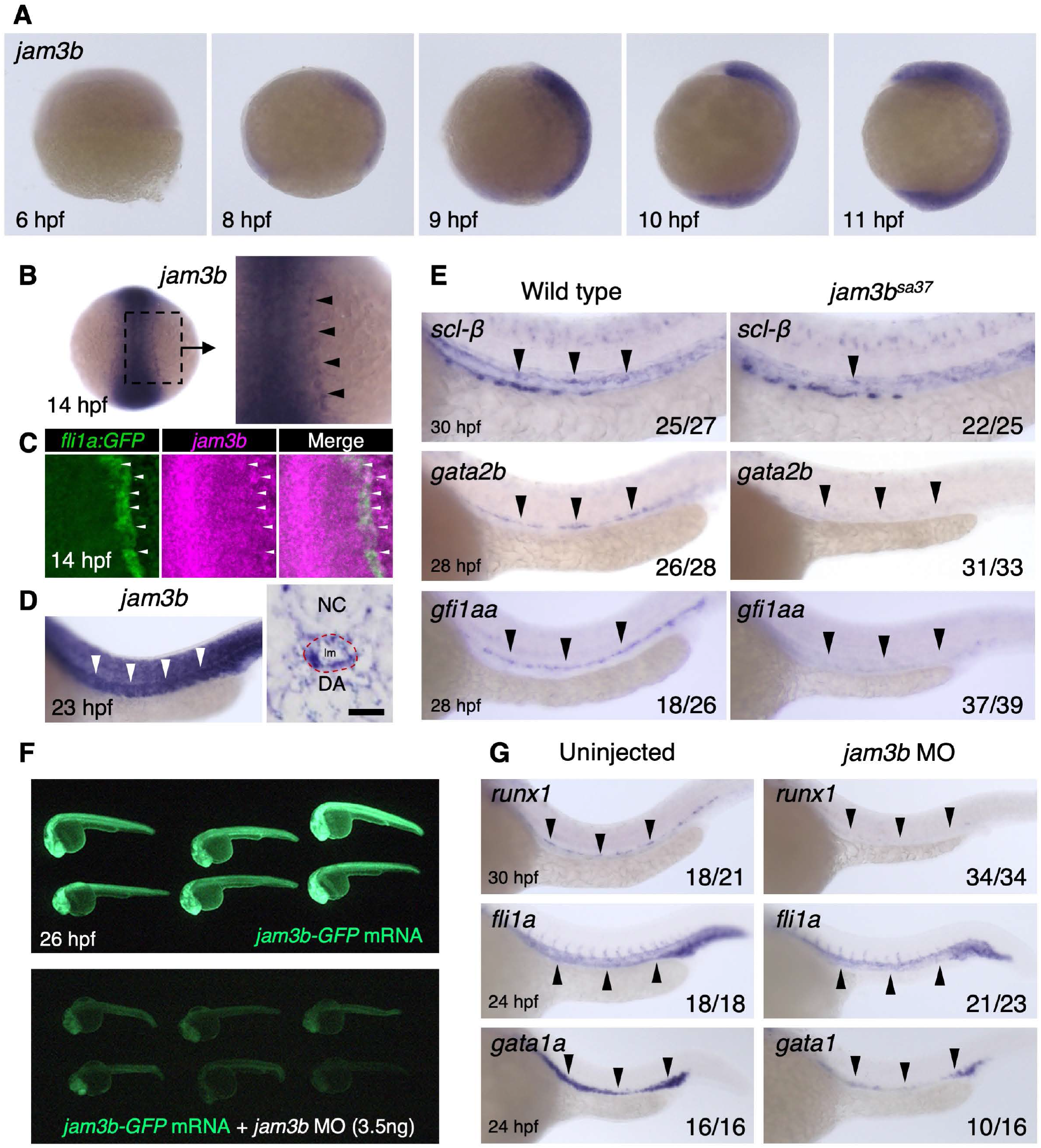
*jam3b* is expressed in the hematopoietic/endothelial tissue. (A) Expression of *jam3b* in WT embryos at 6, 8, 9, 10, and 11 hpf. (B) Expression of *jam3b* in 14 hpf embryos. The right panel shows a high magnification view of the dotted box area in the left panel. Black arrowheads indicate the expression of *jam3b* in the PLPM. (C) Expression of *jam3b* and *fli1a:GFP* in 14 hpf embryos. The expression domain of *jam3b* was partially merged with that of *fli1a:GFP* (white arrowheads). (D) Expression of *jam3b* in 23 hpf embryos. The right panel shows a transverse section of an embryo. White arrowheads in the left panel and a red dotted circle in the right panel indicate the dorsal aorta (DA). NC, notochord; lm, lumen; Bar, 10 µm. (E) Expression of *scl-β*, *gata2b*, and *gfi1aa* in WT or *jam3b^sa37^* embryos. (F) Expression of GFP in embryos injected with *jam3b-GFP* mRNA or co-injected with *jam3b-GFP* mRNA and *jam3b* MO. (G) Expression of *runx1*, *fli1a*, and *gata1a* in embryos uninjected or injected with *jam3b* MO. Black arrowheads in E and G indicate the expression domain of each gene.

**Fig. S2.**
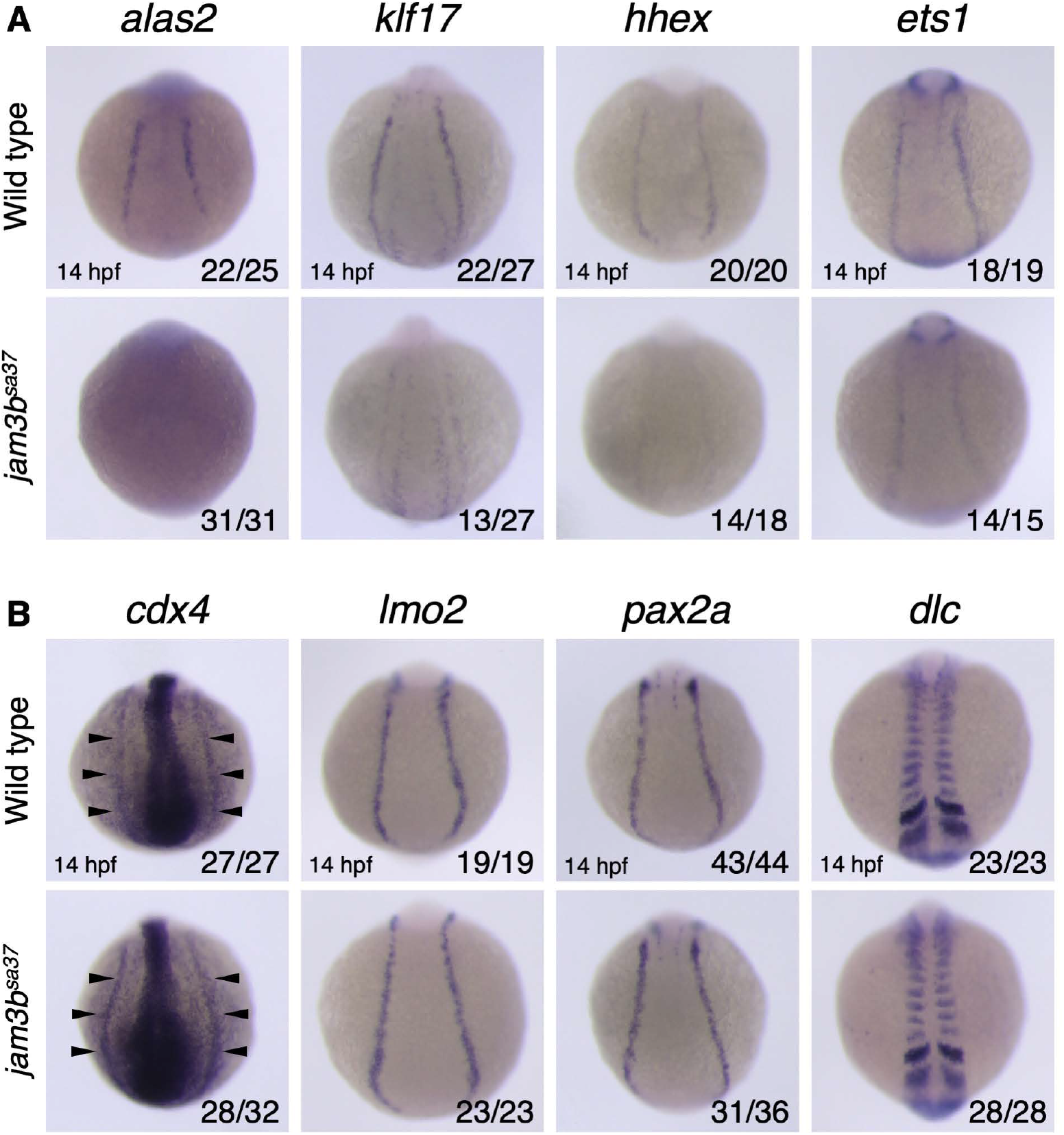
Expression pattern of hematopoietic/endothelial marker genes in *jam3b^sa37^* embryos. (A) Expression of *alas2*, *klf17*, *hhex*, and *ets1* in the PLPM of WT or *jam3b^sa37^* embryos. (B) Expression of *cdx4* and *lmo2* in the PLPM, *pax2a* in the intermediate mesoderm, and *dlc* in the somite of WT or *jam3b^sa37^* embryos. Arrowheads indicate *cdx4* expression in the PLPM.

**Fig. S3.**
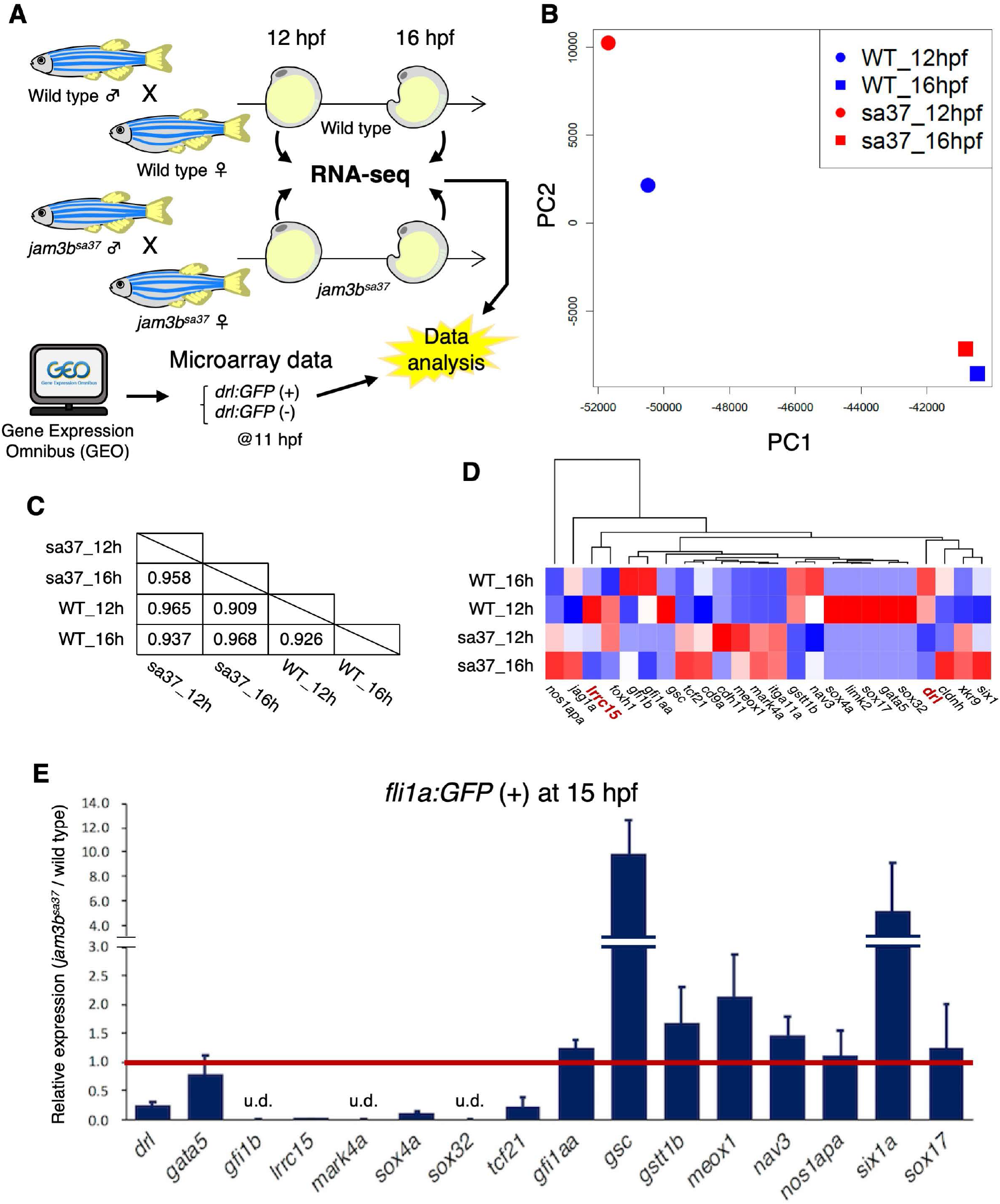
Transcriptome analysis of WT and *jam3b^sa37^* embryos. (A) Experimental outline of RNA-seq analysis in WT and *jam3b^sa37^* embryos. Microarray data of *drl:GFP* (+) and (-) were also used for this analysis. (B) Principal component analysis (PCA) based on the tags per million (TPM) of each sample. (C) Pearson correlation coefficients based on the TPM of each sample. (D) Hierarchical clustering of selected 24 genes based on transcriptome data. (D) Relative expression level of selected genes in *fli1a:GFP*(+) cells from *jam3b^sa37^* embryos. A red bar indicates expression level of each gene in *fli1a:GFP*(+) cells from WT embryos. Error bars, s.d.; u.d., undetected.

**Fig. S4.**
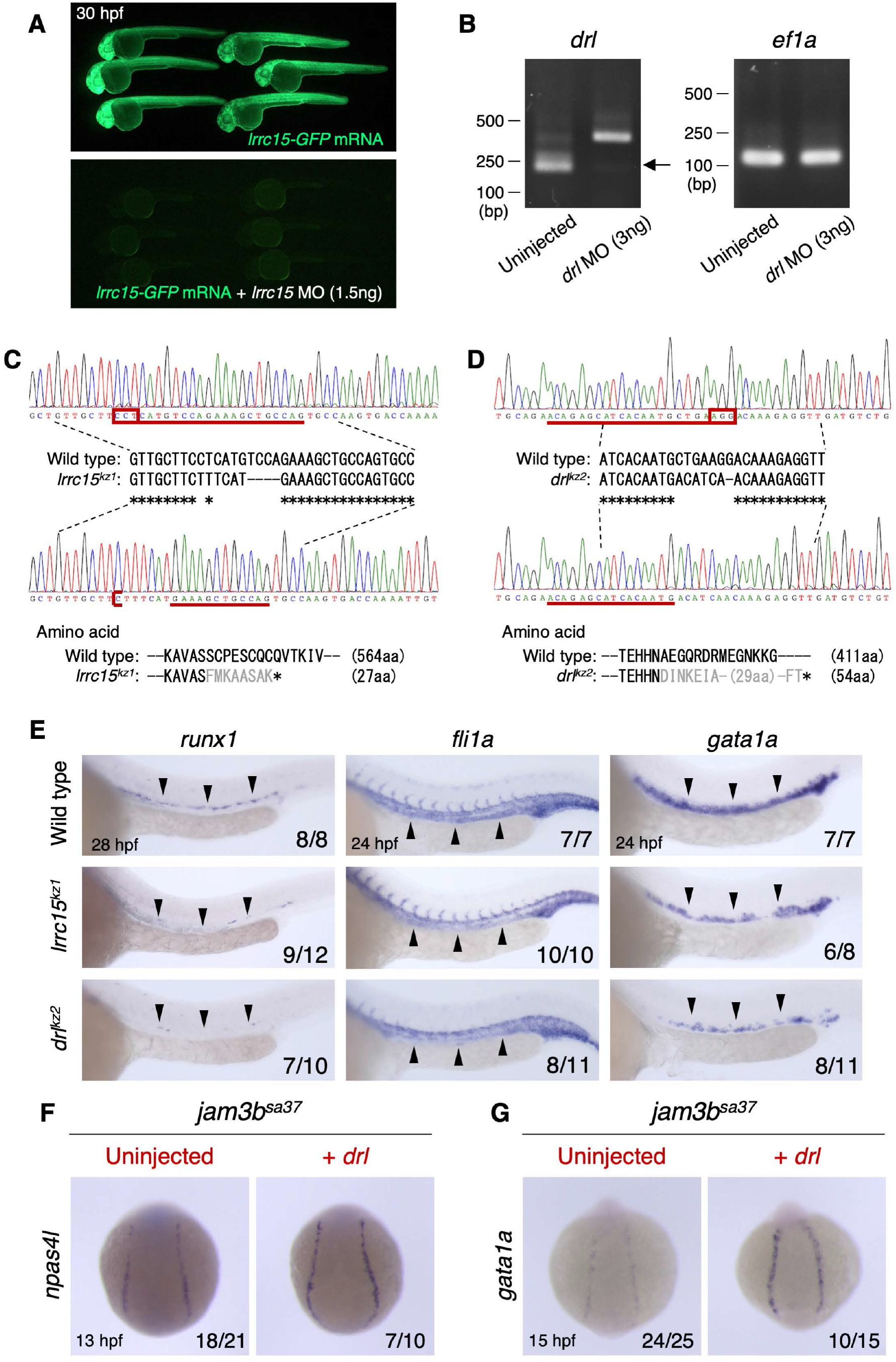
Hemato-vascular development in *lrrc15^kz1^* and *drl^kz2^* embryos. (A) Expression of GFP in embryos injected with *lrrc15-GFP* mRNA or co-injected with *lrrc15-GFP* mRNA and *lrrc15* MO. (B) RT-PCR results of *drl* and *ef1a* in embryos uninjected or injected with *drl* MO. Arrow indicates the expected size of wild type *drl*. (C, D) Mutation in the genomic loci of *lrrc15* and *drl* was verified by sequencing the *lrrc15^kz1^* and *drl^kz2^* lines, respectively. A red box and red line indicate a protospacer adjacent motif (PAM) and the target sequence of gRNA, respectively. Both mutant lines are predicted to have a premature stop codon in exon 2 or 3. (E) Expression of *runx1*, *fli1a*, and *gata1a* in WT, *lrrc15^kz1^*, and *drl^kz2^* embryos. Black arrowheads indicate the expression domain of each gene. (F, G) Expression of *npas4l* and *gata1a* in *jam3b^sa37^* embryos uninjected or injected with *drl* mRNA.

**Fig. S5.**
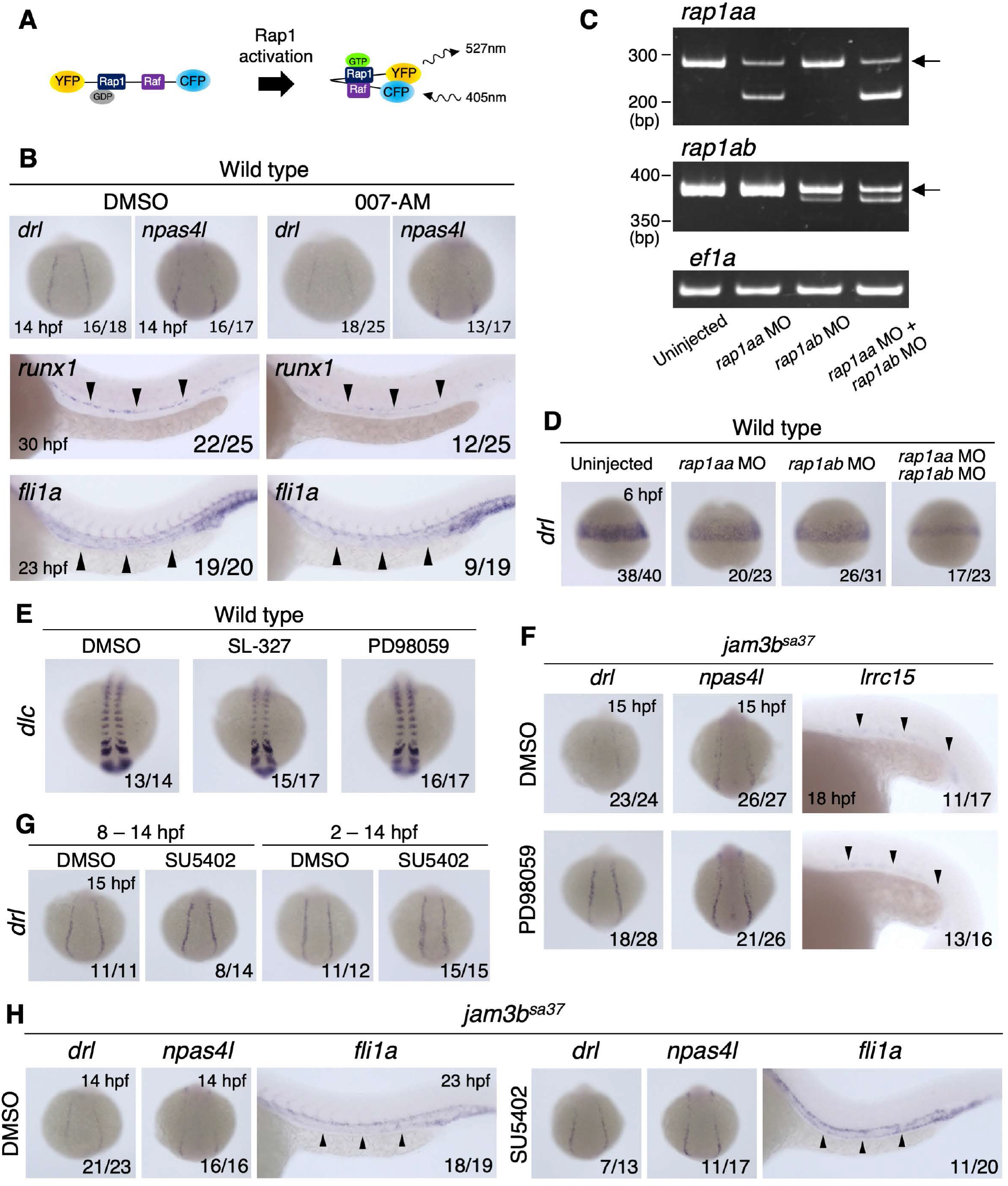
The Rap1a-Erk signaling pathway is regulated by Jam3b. (A) Schematic illustration of Raichu-Rap1 probe. (B) Expression of *drl*, *npas4l*, *runx1*, and *fli1a* in WT embryos treated with DMSO or 007-AM. (C) RT-PCR results from embryos uninjected or injected with *rap1aa* MO, *rap1ab* MO, or both MOs. Expression of *rap1aa*, *rap1ab*, and *ef1a* is shown. Arrows indicate the expected size of wild type *rap1aa* or *rap1ab*. (D) Expression of *drl* in embryos uninjected or injected with *rap1aa* MO, *rap1ab* MO, or both MOs. (E) Expression of *dlc* in WT embryos treated with DMSO, SL-327, or PD98059. (F) Expression of *drl*, *npas4l*, and *lrrc15* in *jam3b^sa37^* embryos treated with DMSO or PD98059. (G) Expression of *drl* in WT embryos treated with DMSO or SU5402 from 8 to 14 hpf (left panels) or 2 to 14 hpf (right panels). (H) Expression of *drl*, *npas4l*, and *fli1a* in *jam3b^sa37^* embryos treated with DMSO or SU5402.

**Fig. S6.**
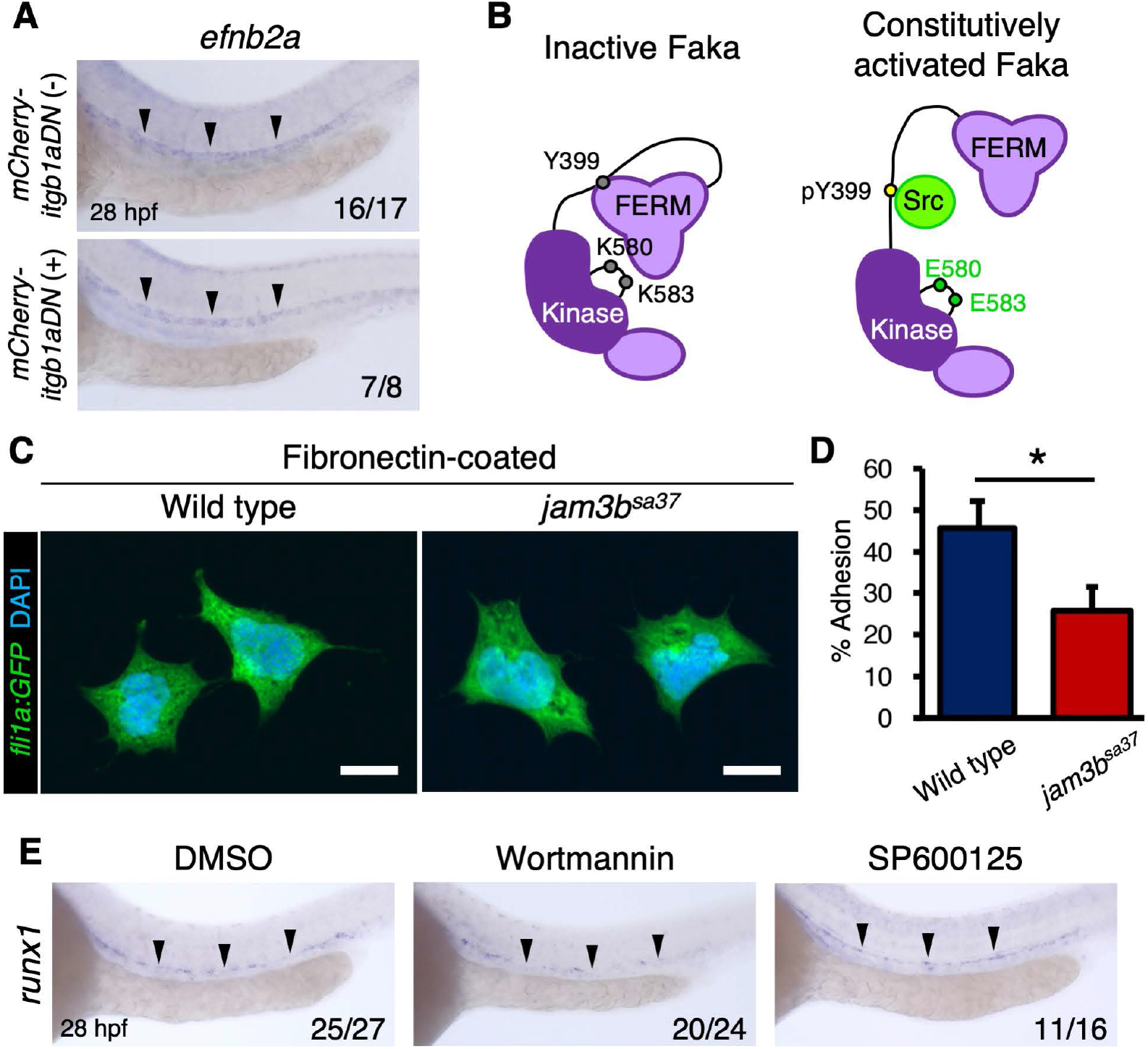
The integrin signaling pathway is regulated by Jam3b. (A) Expression of *efnb2a* in the DA in *mCherry-itgb1aDN* (-) or *mCherry-itgb1aDN* (+) embryos. (B) Schematic illustration of inactive Faka or constitutively activated Faka. FERM, four-point-one, ezrin, radixin, moesin domain. (C) Immunostaining of *fli1a:GFP* (+) cells from WT or *jam3b^sa37^* embryos. GFP (+) cells adherent to a fibronectin-coated glass bottom dish are shown. Bars, 10 µm. (D) Percent adhesion of *fli1a:GFP* (+) cells from WT or *jam3b^sa37^* embryos. Error bars, s.d., \**p* < 0.01. (E) Expression of *runx1* in the DA in WT embryos treated with DMSO, wortmannin, or SP600125.

**Fig. S7.**
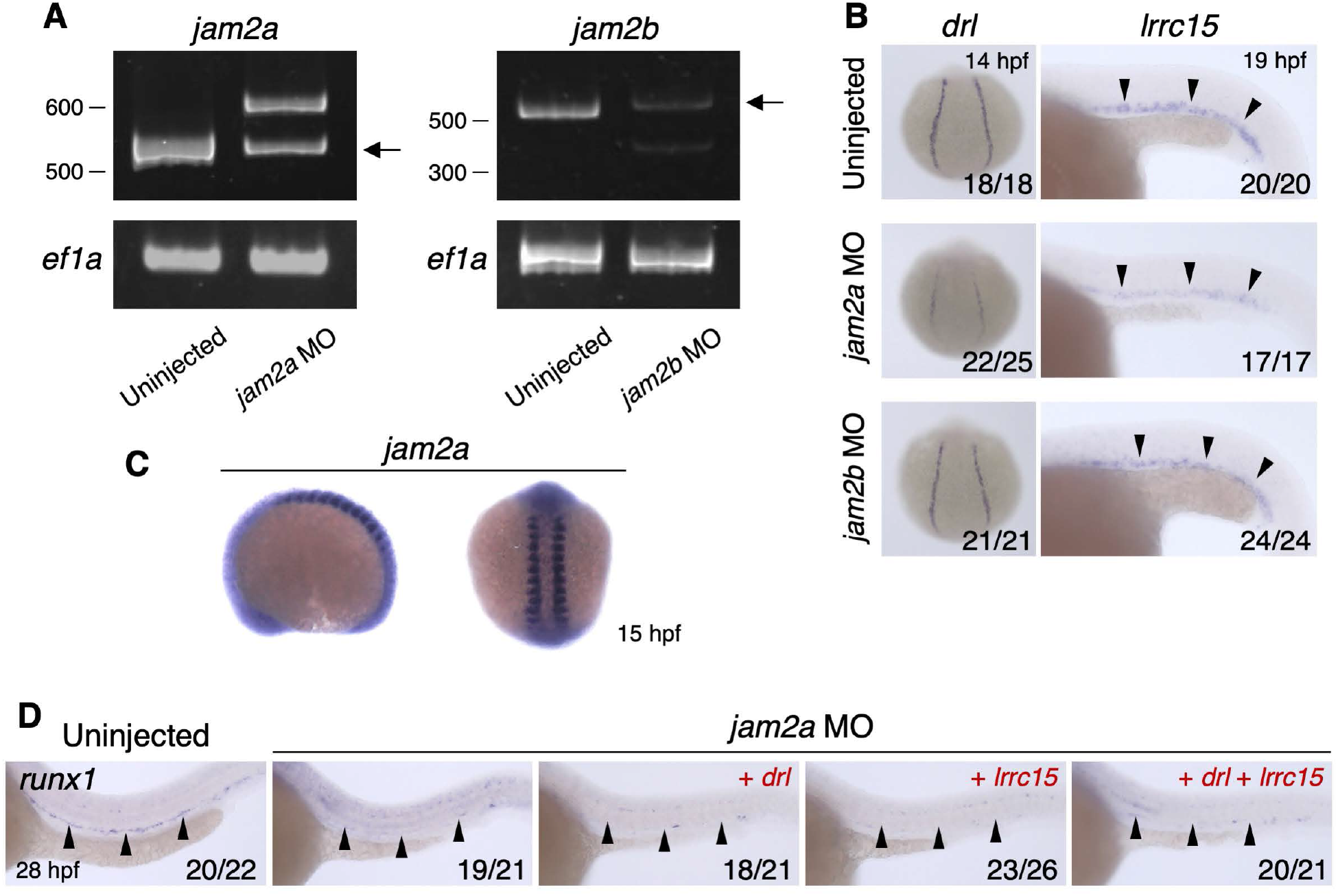
The expression of *drl* and *lrrc15* is Jam2a-dependent. (A) Expression of *jam2a*, *jam2b*, and *ef1a* in embryos uninjected or injected with *jam2a* MO or *jam2b* MO as determined by RT-PCR. Arrows indicate the expected size of wild type *jam2a* or *jam2b*. (B) Expression of *drl* and *lrrc15* in embryos uninjected or injected with *jam2a* MO or *jam2b* MO. (C) Expression of *jam2a* in an 15 hpf embryo. (D) Expression of *runx1* in the DA of uninjected or injected with *jam2a* MO with or without *drl* and *lrrc15* mRNA.

---

**Table S1.** Selected differentially expressed genes between wild type and *jam3b^sa37^* embryos

| Cluster | Gene | Tags per million (TPM) |  |  |  | Fold change<br>*drl*:GFP(+) |
| --- | --- | --- | --- | --- | --- | --- |
|  |  | sa37 | sa37 | WT | WT |  |
|  |  | 12hpf | 16hpf | 12hpf | 16hpf | *drl*:GFP(-) |
| ENSDARG00000005842 | *cd9a* | 0.68 | 1.04 | 0.00 | 0.55 | 2.24 |
| ENSDARG00000021442 | *cdh11* | 0.51 | 0.00 | 0.00 | 0.00 | 4.16 |
| ENSDARG00000090730 | *cldnh* | 1.11 | 8.21 | 0.55 | 4.47 | 2.05 |
| ENSDARG00000069817 | *drl* | 0.85 | 1.73 | 5.49 | 6.97 | 12.09 |
| ENSDARG00000094646 | *foxh1* | 7.52 | 2.49 | 7.61 | 0.86 | 4.35 |
| ENSDARG00000094646 | *gata5* | 0.00 | 0.00 | 1.25 | 0.00 | 12.87 |
| ENSDARG00000094646 | *gfi1aa* | 1.28 | 0.58 | 2.12 | 3.79 | 2.50 |
| ENSDARG00000094646 | *gfi1b* | 0.60 | 1.21 | 0.00 | 2.75 | 2.34 |
| ENSDARG00000094646 | *gsc* | 0.00 | 0.00 | 3.21 | 0.00 | 3.45 |
| ENSDARG00000094646 | *gstt1b* | 0.00 | 0.00 | 0.78 | 0.86 | 3.03 |
| ENSDARG00000094646 | *itga11a* | 0.60 | 0.58 | 0.00 | 0.00 | 3.58 |
| ENSDARG00000094646 | *jag1a* | 7.94 | 11.79 | 3.53 | 9.18 | 2.66 |
| ENSDARG00000094646 | *limk2* | 0.00 | 0.00 | 0.71 | 0.00 | 2.23 |
| ENSDARG00000094646 | *lrrc15* | 2.05 | 0.00 | 6.59 | 1.28 | 2.85 |
| ENSDARG00000094646 | *mark4a* | 0.51 | 0.58 | 0.00 | 0.00 | 2.45 |
| ENSDARG00000094646 | *meox1* | 0.94 | 0.52 | 0.00 | 0.00 | 3.96 |
| ENSDARG00000062427 | *nav3* | 0.00 | 0.52 | 0.55 | 0.98 | 2.49 |
| ENSDARG00000062427 | *nos1apa* | 20.33 | 33.18 | 4.16 | 1.35 | 2.33 |
| ENSDARG00000008029 | *six1a* | 3.08 | 5.61 | 1.33 | 3.12 | 3.63 |
| ENSDARG00000090408 | *sox17* | 0.00 | 0.00 | 0.86 | 0.00 | 9.43 |
| ENSDARG00000090408 | *sox32* | 0.00 | 0.00 | 1.65 | 0.00 | 10.21 |
| ENSDARG00000090408 | *sox4a* | 0.00 | 0.00 | 0.55 | 0.00 | 2.13 |
| ENSDARG00000090408 | *tcf21* | 0.77 | 1.16 | 0.00 | 0.00 | 2.30 |
| ENSDARG00000117533 | *xkr9* | 4.70 | 5.26 | 0.86 | 1.65 | 4.94 |

**Table S2.**
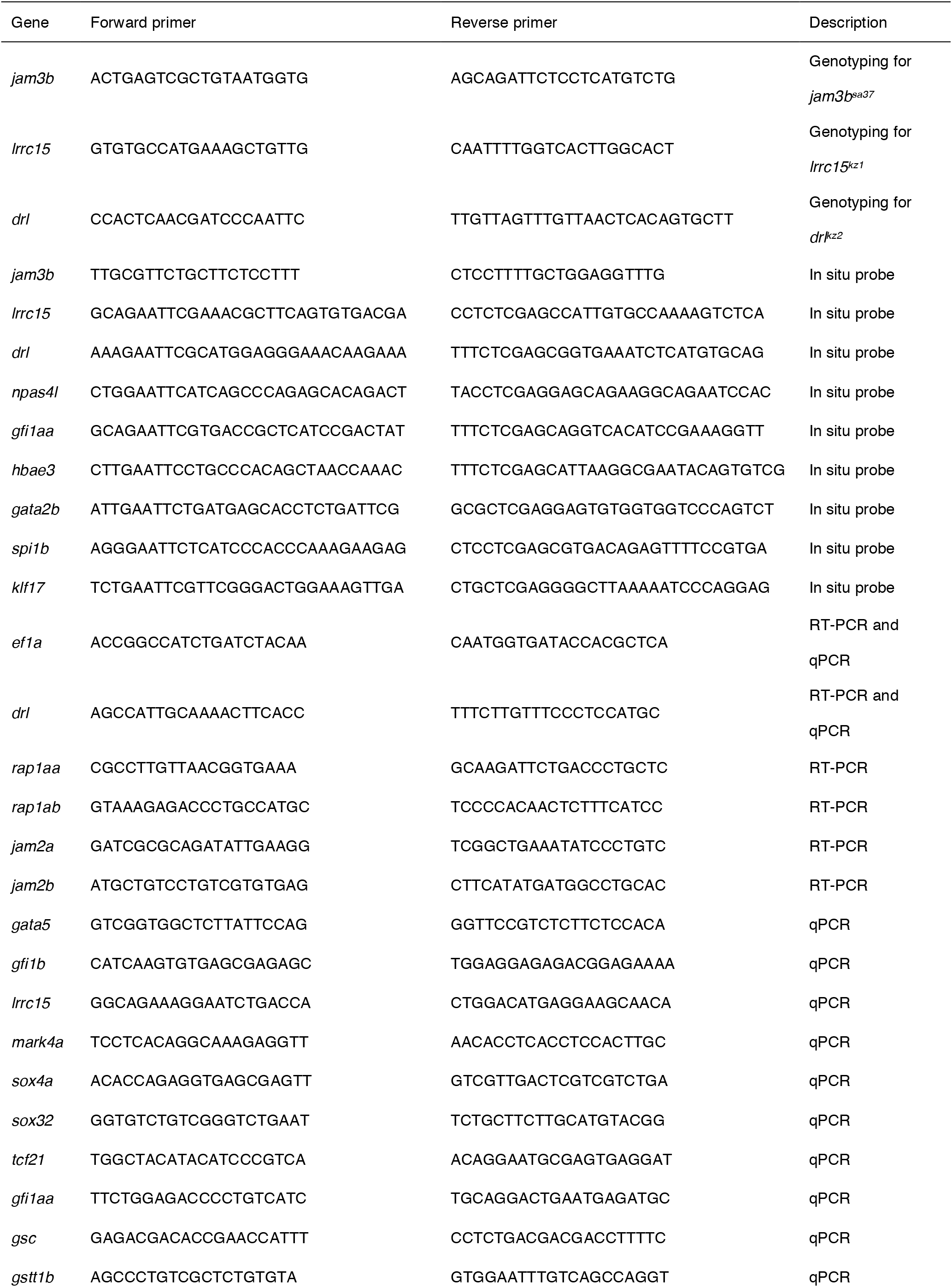
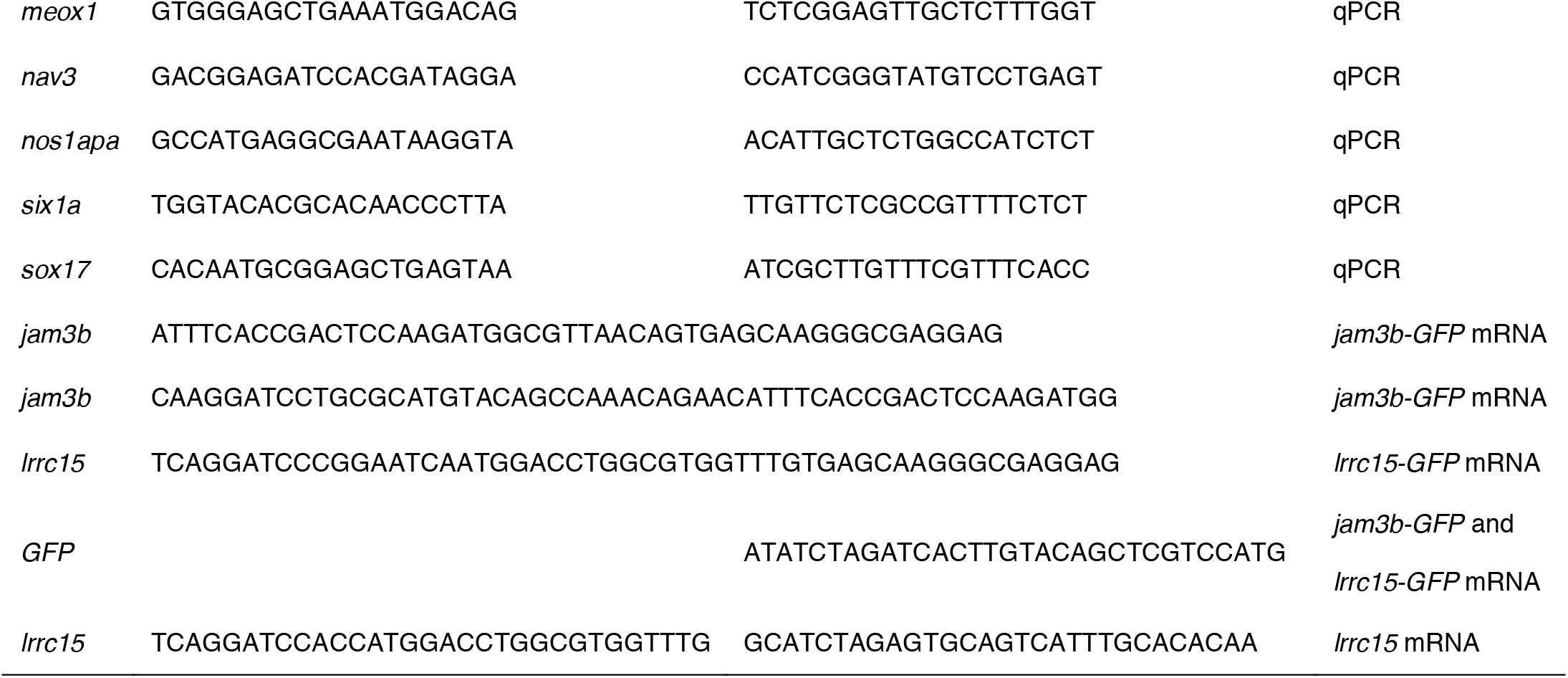
Sequences for primers

